# High-Resolution Dissection of Conducive Reprogramming Trajectory to Ground State Pluripotency

**DOI:** 10.1101/184135

**Authors:** Asaf Zviran, Nofar Mor, Yoach Rais, Hila Gingold, Shani Peles, Elad Chomsky, Sergey Viukov, Jason D. Buenrostro, Leehee Weinberger, Yair S. Manor, Vladislav Krupalnik, Mirie Zerbib, Hadas Hezroni, Diego Adhemar Jaitin, David Larastiaso, Shlomit Gilad, Sima Benjamin, Awni Mousa, Muneef Ayyash, Daoud Sheban, Jonathan Bayerl, Alejandro Aguilera Castrejon, Rada Massarwa, Itay Maza, Suhair Hanna, Ido Amit, Yonatan Stelzer, Igor Ulitsky, William J. Greenleaf, Yitzhak Pilpel, Noa Novershtern, Jacob H. Hanna

**Affiliations:** Department of Molecular Genetics, Weizmann Institute of Science, Rehovot 7610001, Israel.; Department of Neurobiology, Wise Faculty of Life Sciences and Sagol School of Neuroscience, Tel Aviv University, Tel Aviv 69978, Israel.; Department of Biological Regulation, Weizmann Institute of Science, Rehovot 7610001, Israel.; Department of Computer Science, Weizmann Institute of Science, Rehovot 7610001, Israel.; Broad Institute of MIT and Harvard, Cambridge, MA 02142, USA; Harvard Society of Fellows, Harvard University, Cambridge, MA 02138, USA; Department of Immunology, Weizmann Institute of Science, Rehovot 76100, Israel.; The Israel National Center for Personalized Medicine (INCPM), Weizmann Institute of Science, Rehovot 76100, Israel.; Department of Gastroenterology, Rambam Health Care Campus & Bruce Rappaport School of Medicine, Technion Institute of Technology, Haifa, Israel; Department of Pediatrics and the Pediatric Immunology Unit, Rambam Health Care Campus & Bruce Rappaport School of Medicine, Technion Institute of Technology, Haifa, Israel; Department of Molecular and Cell Biology, Weizmann Institute of Science, Rehovot 76100, Israel.; Department of Applied Physics, Stanford University, Palo Alto, CA, USA; Chan Zuckerberg Biohub, San Francisco, CA 94158, USA

## Abstract

The ability to reprogram somatic cells into induced pluripotent stem cells (iPSCs) with four transcription factors Oct4, Sox2, Klf4 and cMyc (abbreviated as OSKM)^1^ has provoked interest to define the molecular characteristics of this process^2-7^. Despite important progress, the dynamics of epigenetic reprogramming at high resolution in correctly reprogrammed iPSCs and throughout the entire process remain largely undefined. This gap in understanding results from the inefficiency of conventional reprogramming methods coupled with the difficulty of prospectively isolating the rare cells that eventually correctly reprogram into iPSCs. Here we characterize cell fate conversion from fibroblast to iPSC using a highly efficient deterministic murine reprogramming system engineered through optimized inhibition of Gatad2a-Mbd3/NuRD repressive sub-complex. This comprehensive characterization provides single-day resolution of dynamic changes in levels of gene expression, chromatin modifications, TF binding, DNA accessibility and DNA methylation. The integrative analysis identified two transcriptional modules that dominate successful reprogramming. One consists of genes whose transcription is regulated by on/off <u>e</u>pigenetic <u>s</u>witching of modifications in their <u>p</u>romoters (abbreviated as ESPGs), and the second consists of genes with <u>p</u>romoters in a <u>c</u>onstitutively <u>a</u>ctive chromatin state, but a dynamic expression pattern (abbreviated as CAPGs). ESPGs are mainly regulated by OSK, rather than Myc, and are enriched for cell fate determinants and pluripotency factors. CAPGs are predominantly regulated by Myc, and are enriched for cell biosynthetic regulatory functions. We used the ESPG module to study the identity and temporal occurrence of activating and repressing epigenetic switching during reprogramming. Removal of repressive chromatin modifications precedes chromatin opening and binding of RNA polymerase II at enhancers and promoters, and the opposite dynamics occur during repression of enhancers and promoters. Genome wide DNA methylation analysis demonstrated that de novo DNA methylation is not required for highly efficient conducive iPSC reprogramming, and identified a group of super-enhancers targeted by OSK, whose early demethylation marks commitment to a successful reprogramming trajectory also in inefficient conventional reprogramming systems. CAPGs are distinctively regulated by multiple synergystic ways: 1) Myc activity, delivered either endogenously or exogenously, dominates CAPG expression changes and is indispensable for induction of pluripotency in somatic cells; 2) A change in tRNA codon usage which is specific to CAPGs, but not ESPGs, and favors their translation. In summary, our unbiased high-resolution mapping of epigenetic changes on somatic cells that are committed to undergo successful reprogramming reveals interleaved epigenetic and biosynthetic reconfigurations that rapidly commission and propel conducive reprogramming toward naïve pluripotency.

## Main Text

Elegant previous epigenetic mapping studies on iPSC reprogramming were conducted on inefficient and a-synchronized systems^6,8^ undergoing prolonged and protracted reprogramming course (e.g. over 20 days and multiple passages^3,9^) or by sorting subpopulations and “pre-iPSCs” at specific time points, most of which do not progress to become iPSCs (e.g. SSEA1+ cells)^3,5,9^. However, this low-efficiency and heterogeneity has limited genome-wide analysis of well-characterized, relatively homogeneous populations of cells that rapidly and successfully complete this process. Further, none of these studies were carried in naïve 2i/LIF (2 inhibitors for MEK and GSK3 pathways + LIF cytokine) ground state pluripotency conditions, which are known to endow unique epigenetic configuration and regulation of resultant pluripotent cells^10,11^.

Multiple studies have recently reported alternative reprogramming methods that can achieve deterministic iPSC reprogramming from mouse somatic cells, reaching up to 95-100% iPSC reprogramming efficiency within 8-10 days following OSKM induction. Transient pre-induction of murine B cells with C/EBPa followed by OSKM induction in naïve 2i/LIF conditions yields 85-100% Oct4-GFP reporter within 8 days^12^. Our group has demonstrated that optimized hypomorphic depletion of Mbd3, representing a core member of the Mbd3/Chd4/NuRD co-repressor complex, followed by supplementing 2i/LIF/KSR at day 3, results in (near-)deterministic and more synchronized reprogramming in mouse cells within 8 days^13^, ^14^. Notably, independent single cell analysis by CyTOF has confirmed the efficiency and ESC-like molecular authenticity of iPSCs obtained from Mbd3^flox/-^ system following only 8 days of reprogramming^14^. We have also found that complete inhibition of Gatad2a (also known as P66a), a NuRD specific subunit, does not compromise somatic cell proliferation as previously seen upon complete Mbd3 protein elimination, and yet disrupts Mbd3/NuRD repressive activity on the pluripotent circuitry and yields 90-100% highly-efficient reprogramming within 8 days as similarly observed in Mbd3 hypomorphic donor somatic cells (Extended Data Fig. 1 and Mor N. et al.^15^ *Under final preparation*). Such systems now enable high-resolution temporal dissection of epigenetic dynamics underlying conducive naïve iPSC formation, while simultaneously eliminating ‘noise’ from heterogeneous populations that fail to correctly embark or successfully complete the reprogramming process.

With this unique opportunity to map epigenetics of reprogramming towards ground state naïve pluripotency in highly efficient and homogenous systems, without cell passaging or sorting for subpopulations, we provide herein comprehensive characterization during the entire 8-day course of fibroblast reprogramming, at a single day resolution, by measuring transcription, chromatin modifications, DNA accessibility, DNA methylation and binding of transcription factors. These data provide insights into key questions underlying successful somatic cell reprogramming to naive pluripotency: What is the temporal progression of epigenetic reprogramming that occurs during cellular reprogramming? How do epigenetic changes control the induction and repression of gene modules during successful reprogramming? Does DNA demethylation occur in waves, or is it a continuous process? Is there an early epigenetic indicator for successful reprogramming trajectory?

### Dissecting conducive reprogramming in two independent NuRD-deficient systems

We utilized the high efficiency of Gatad2a-Mbd3/NuRD optimally depleted reprogrammable systems, and analyzed reprogramming of two independently generated Mbd3^f/-^ and Gatad2a^-/-^ secondary mouse embryonic fibroblast (MEF) clonal systems, carrying a doxycycline (DOX) inducible STEMCCA-OKSM lentiviral reprogramming vector as previously described^13^ (Extended Data Figure 1). Briefly, Mbd3^f/-^, Gatad2a^-/-^ or wild-type (WT) MEFs that carry DOX-inducible OKSM cassette were used (**Methods**), and iPSC reprogramming was initiated by addition of DOX in FBS/LIF medium in 5% pO_2_ conditions (Fig. 1a). On day 3.5 after DOX initiation, medium was changed to LIF/KSR-based with the addition of 2i. For all mouse iPSC reprogramming experiments, irradiated human foreskin fibroblasts were used as feeder cells, as any sequencing input originating from the use of human feeder cells cannot be aligned to the mouse genome and is therefore omitted from the analysis. Cells were harvested every 24 hours until day 8, and processed for library preparation followed by high-throughput sequencing. Mbd3^f/-^, Gatad2a^-/-^ and WT established iPSC line (after 3 passages or more), and Mbd3^f/-^ or WT V6.5 mouse ESCs were used as positive controls. Two independent WT MEF secondary reprogramming systems (WT-1 and WT-2) were used as additional controls, where WT-2 is an isogenic genetically matched cell line to the Gatad2a^-/-^ cells used herein (Extended Data Fig. 1). The use of independent Mbd3-and Gatad2a-depleted highly efficient systems with different OKSM transgene integrations, excludes cell line specific signatures. We sequenced various measurements in up to eleven time points, resulting in overall 212 libraries from the NuRD depleted systems and 21 libraries from control WT-1/2 systems (**Supplementary Table S1**). The libraries span transcriptome (RNA-Seq, small RNA-seq), chromatin modifications (ChIP-seq for H3K27ac, H3K27me3, H3K4me1, H3K4me3, H3K36me3, H3K9me2, H3K9me3), DNA methylation (reduced representation bisulfite sequencing (RRBS), whole-genome bisulfite sequencing (WGBS)), chromatin accessibility (ATAC-seq), and transcription factor binding (Oct4, Sox2, Klf4, c-Myc and RNA-PolII ChIP-seq). Overall, we aligned 12.12 billion reads, with an average of 50M aligned reads per sample (**Supplementary Table S1**).

**Fig. 1:**
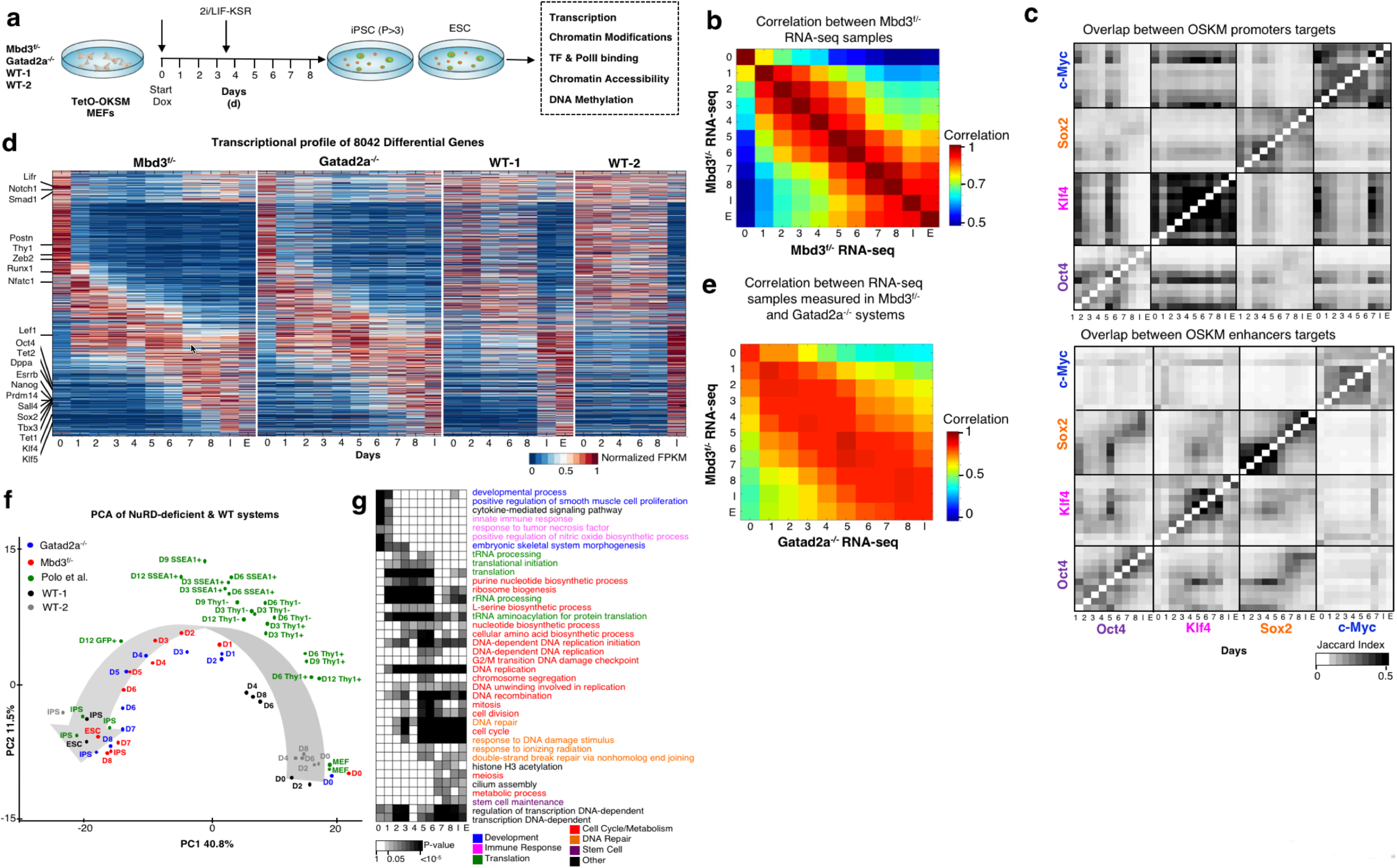
Continuous and coordinated progression of conducive reprogramming in two independent NuRD-deficient systems. a. Experimental Flow. Mbd3^f/-^ and Gatad2a^-/-^ mouse fibroblasts were induced to OKSM reprogramming by Doxycycline. Samples were harvested every 24h during entire 8-day course of reprogramming, without sorting or passaging. Established iPSCs and ESCs were taken as control. Transcription, chromatin modifications, TF binding, DNA accessibility and DNA methylation were measured (see Table S1 for details). b. Spearman correlation between expression profiles of Mbd3^f/-^ system. Calculated over all differential genes (n=8,042), showing an average correlation of R=0.93 between consecutive samples. c. Overlap between targets of OSKM in promoters (top) and enhancers (bottom). Pixel shade indicates Jaccard Index, which is defined as |X^Y|/|XvY|. All transcription factors show high similarity between consecutive samples. Oct4 and Sox2 change their target enhancers throughout the process, and to less extent, their promoter targets. Enhancer targets of Oct4 and Sox2 highly overlap. Klf4 targets overlap with those of Oct4 and Sox2 mostly in intermediate reprogramming stages. d. Global transcriptional pattern of 8,042 (PolyA+) differential genes (FC>4 & maximal FPKM value>1), sorted by their temporal pattern in Mbd3^f/-^ system (The same gene order was applied for the other reprogramming systems). Heatmap represents unit-transformation of FPKM values (see Methods). f. PCA analysis of all samples, alongside samples from previous publications^5^. PCA was calculated on the same set of genes and normalization as in panel (d). g. GO categories enriched among the genes that are active in each day. Gene is defined to be active in samples where RPKM is above 0.5 of the gene max value. P-values were calculated with Fisher exact test, and FDR corrected. Categories with corrected p-value<0.01 in at least two-time points are presented. Gray Shades represent FDR corrected p-values

RNA-seq samples were highly reproducible with average correlation of R=0.93 between consecutive samples (Fig. 1b). The chromatin modification data were also reproducible with correlation above 0.5 between consecutive samples (Extended Data Fig. 2a). TF ChIP-seq samples showed high overlap between consecutive samples (Jaccard index >0.3, Fig. 1c). In addition, iPSC and ESC samples were compared to previously published data^16-18^, and showed high consistency with the previously established peaks (Fisher exact test p<10^-4^, Extended Data Fig. 2b), thus confirming the quality of ChIP-seq and other datasets generated and analyzed herein.

### Conducive reprogramming is accompanied by continuous transcriptional changes

Gene expression profiles in Mbd3^f/-^ and Gatad2a^-/-^ systems were highly similar (Fig. 1d-e, Extended Data Fig. 2c, **Supplementary Tables S2, S3**), despite the independent OKSM lentiviral integrations and different genetic background (average R=0.88), further supporting the conclusion that both optimally NuRD-depleted systems induced rapid and efficient reprogramming. To further evaluate the kinetics of the two systems, we compared them to two WT secondary reprogramming systems (WT-2 series is isogenic to Gatad2a^-/-^ series), and to previously published dataset which mapped iPSC reprogramming from WT fibroblasts and isolated different intermediate populations based on surface marker expression (Thy1 and SSEA1 markers)^5^. Principal component analysis (PCA) mapped the RNA-seq samples in a trajectory that reflects the progression of reprogramming from MEF to IPS/ES (Fig. 1f). MEF samples (i.e. day 0) of all systems are clustered together in close proximity to WT samples measured in days 2, 4, 6 and 8, emphasizing the fast kinetics of NuRD-depleted systems compared to WT systems, and that the starting NuRD-depleted MEFs are not transcriptionally different from WT MEFs. IPS and ES samples of all systems are also clustered together, along with Mbd3^f/-^ and Gatad2a^-/-^ samples taken at day 8 after DOX induction, reiterating complete induction of pluripotency in nearly all donor somatic cells. Mbd3^f/-^ and Gatad2a^-/-^ samples from the same time points are closely mapped in both dimensions (Fig. 1f). Importantly, although previously published data^5^ measured in WT MEF and IPS is clustered together with our corresponding samples, all other samples, which were measured from sorted cells undergoing reprogramming, are positioned in clusters based on the marker used for sorting and not according to the reprogramming day (Fig. 1f - compare Polo et al.^5^ data points to Mbd3^flox/-^ and Gatad2a^-/-^). In addition, Thy1+, Thy1-and SSEA1+ WT per-iPSC samples do not cluster with any of the samples from Mbd3^f/-^ or Gatad2a^-/-^ systems beyond day 3 of reprogramming (PC1), consistent with the notion that reprogramming measurements on inefficient stochastic reprogramming systems focus on early time points and infer data on populations most of which are “trapped”^5,9,19^ and not necessarily proceeding toward becoming iPSCs. In summary, the above analysis validates the high quality of the systems used herein and data generated from them, and underscores the relevance of studying continuous non-saltatory trajectory of highly-efficient successful iPSC reprogramming.

To characterize transcriptional trajectory during deterministic reprogramming, polyA+ RNA-seq measuring mRNAs and long noncoding RNAs (lncRNAs), and small RNA-seq measuring microRNAs (miRNAs) were applied. Previous analysis on stochastic reprogramming systems have indicated two waves of major transcriptional changes, one at the beginning and the other at the end of iPSC reprogramming, while in between there are very minor changes in transcriptional patterns^5^. We set out to test whether highly efficient reprogramming systems will show a similar pattern. 8,705 genes (of which 8,042 are polyA+) were identified as differentially expressed along the Mbd3^f/-^ MEF to naïve iPSC 8-day reprogramming course (**Supplementary Table S3**). These genes show a sequential activity, and can be sorted according to their expression temporal pattern, showing a continuous dynamic transition from the somatic program to the pluripotent one. Three major expression shifts are observed during the continuous dynamic transition (Fig. 1g, Extended Data Fig. 2c): First, a large group of genes which are active in MEFs are down regulated as early as day 1. The second is a transient activation of genes between days 1 and 4. Finally, there is a gradual establishment of iPS/ES signature starting at day 5. Functional enrichment analysis in a single day resolution (Fig. 1g) characterized these changes: genes which are active in MEF and downregulated after DOX induction are enriched for somatic program processes (e.g. developmental process, skeletal system, cytokine-mediated signaling) ^4,5,20^. Genes induced between days 1 and 6 are enriched for processes related to biosynthetic pathways (DNA synthesis, DNA replication, purine biosynthesis, translation, ribosome biogenesis). These processes are followed by induction of genes enriched for epigenetic remodeling and DNA repair processes (e.g. chromosome segregation, DNA repair, DNA recombination, response to DNA damage). Finally, at day 6 there is a prominent induction of pluripotency maintenance master regulators including Nanog, Esrrb, Tbx3, Sall4, Prdm14 (Extended Data Fig. 3a)^4,5,20,21^. In summary, at the transcriptional level, during conducive reprogramming trajectory, somatic cell repression and pluripotency gene reactivation associated changes do not occur simultaneously and are separated in time. However, many other changes related to cellular adaptation occur in between, thus rendering global transcriptional changes rather continuous and not confined only to early and late stages of iPSC reprogramming as suggested before^5,9^. An expanded lncRNA (**Methods**) and miRNAs^18,22^ mapping showed continuous patterns of activation or repression during the reprogramming process, as seen with protein coding genes (Extended Data Fig. 3b, c; **Supplementary Table 4**).

### Dynamic OSK binding governs conducive iPSC formation

Along with the change in gene transcription, we identified 40,174 enhancers with a dramatic change in activity, which is consistent between the two Gatad2a/Mbd3-NuRD depletion approaches used (Extended Data Figs. 4,5 **and methods**). The changes in gene expression and enhancer activity are first triggered by the induction of OSKM factors^23-25^ and thus we next characterized the binding of OSKM. We observed dynamic binding pattern for OSK in enhancers (Fig. 1c), which is changing during reprogramming from an early pattern to late pluripotency related pattern (Fig. 2a-c). cMyc has a strong preference to bind promoters over enhancers during reprogramming (Fig. 2b), similar to what can be observed during ESC maintenance^25-28^. Inspecting the binding co-localization of OSKM shows a clear difference between promoters and enhancers. Oct4-binding enhancers overlap with Sox2 and to a lesser extent with Klf4 targets throughout the process (Fig. 2d). The average probability to see co-localization of Oct4 and Sox2 in enhancers is 0.77 while in promoters it decreases to 0.44. However, the probability of Oct4 and Klf4 co-localization in promoters is higher by ~2 fold compared to enhancers (Fig. 2d). Differences between binding of enhancers and promoters are also apparent at the DNA motif level (Extended Data Fig. 6). While OSK-binding promoters are enriched mainly for temporally stable OSK binding motifs, OSK-binding enhancers are enriched for many additional and temporally varying binding motif patterns (Extended Data Fig. 6). For example, Sox2 target enhancers are enriched for Oct4-Sox17, Ap1 and NF-E2 motifs (early), Ap2alpha/gamma, E2f1 and Sp1 motifs (intermediate), and Esrrb, Nr5a2 and Tcf4 motifs (late). This change in motif preference may indicate a change in the collaborative binding of OSK during reprogramming^20^, and may be responsible for the more dramatic changes in enhancer activity (-5-fold increase in total differential enhancers (40,174) in comparison to differential gene promoters (8,042) underscores the dramatic magnitude of enhancer reprogramming).

**Fig. 2:**
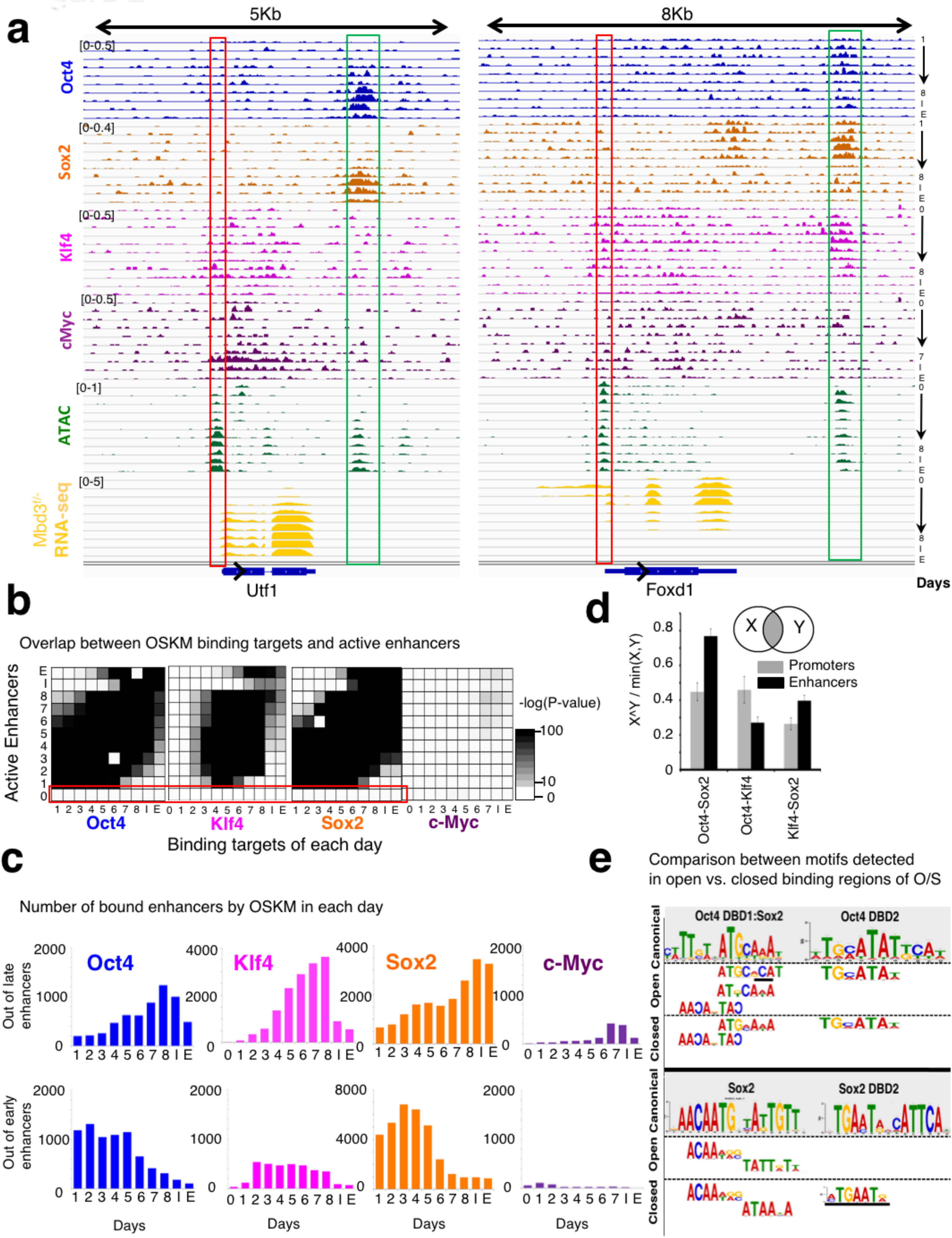
Stage-specific binding preference and collaboration of OSKM. a. ChIP-seq landscape of two gene examples, showing binding of Oct4, Sox2, Klf4 and c-Myc, alongside ATAC-seq and RNA-seq. Promoters are marked in red, enhancers are marked in green. Note the dynamic nature of Oct4 and Sox2 binding. Signals are normalized to sample size (RPM), and IGV data range is indicated on top left corner of the signal. b. Overlap between binding of OSKM and active enhancers, in each day of reprogramming. Enhancer is defined as active in a specific day if its ATAC-seq z-score is above 1.5 STD in that day. Gray shades indicate Fisher exact test p-value for overlap between compared samples. Note that OSKM do not bind the enhancers that are active in MEF (D0, marked in red); these enhancers are not significantly bound by OSKM at any day during reprogramming. c. Number of enhancers bound by each of OSKM factors in each day of reprogramming. Upper row: out of enhancers that are bound by the factor in late stages (day8, IPS, ESC). Bottom row: out of enhancers that are bound by the factor in early stages (day1-day3). d. Probability to observe co-localized binding of transcription factors in promoters (gray) and enhancers (black). Calculated in days 0,1,8 and IPS (Error bars indicate S.E.M). e. Motifs found in “closed” vs. “open” binding targets of the indicated transcription factor. Accessibility of targets was calculated based on ATAC-seq (See Methods). Motifs found in OSK binding targets calculated in Mbd3^f/-^ day1. Motifs that are different between open and closed binding targets are marked in black line. Complementary motifs to canonical motif appear in reverse order.

We next asked if OSK are directly responsible for the repression of the somatic program. When inspecting enhancers that are active in MEF and repressed already at day 1, we observed that they are not significantly bound by OSK at any stage (not even when OSK are already expressed at day 1) (Fig. 2b). This observation is different from that reported by Chronis et al.^20^, who observed predominant OSK binding on MEF open enhancers at day 2 of the process. This difference is likely due to the usage of a system with very low successful iPSC efficiency (less than 0.1%^20,29^), where most donor cells fail to correctly engage in conducive reprogramming. Our observations suggest that different regulators may mediate the repression of MEF-enhancers during successful reprogramming. Indeed, MEF-enhancers are enriched for binding motif of Runx1, Tead, Nf1 and Erg (p-val <10^-70^, Extended Data Fig. 7a), and to much lower extent to Oct4 or Sox2 (p-val =10^-6^). This begins to change from day 1, where active enhancers are significantly enriched (p<10^-250^) for Oct4 and Sox2 binding motif. This includes late stage enhancers that are enriched (p<10^-50^) for other pluripotent transcription factors such as Esrrb and Prdm14 which are upregulated during the process (Extended Data Fig. 7b). It has been previously shown that Oct4 and Sox2 are pioneer factors which have the ability to bind closed chromatin and to activate new regulatory elements^24,30,31^. Specifically, 70% of the enhancers bound by OSK after 48h of reprogramming are in a closed chromatin state in human fibroblasts^24^. In the highly efficient mouse system used here, out of the 4,858 enhancers that are bound by OSK on day 1, 74% were in a closed state (no mark) in MEF and 4% were repressed by either H3K9me2 or H3K27me3. Further, when we examined the binding motifs abundant in closed vs. opened binding sites of OSK in day1 of Mbd3^f/-^ system, the canonical TF binding motifs of OSK were detected after 1 day of DOX induction in Mbd3^flox/-^ cells in both closed and open regions (Fig. 2e, Extended Data Fig. 7c). The latter can be consistent with a pioneer TF activity for OSK^24^ in conducive mouse reprogramming.

### Early enhancer demethylation marks commissioning of conducive reprogramming

The reprogramming of cellular fate involves not only changes in transcription and binding of transcription factors, but also changes in DNA methylation^3-5^. It has been shown that DNA methylation is eventually reduced during cellular reprogramming in pluripotency genes^5,13,32^, however up-regulation of DNA methylation over lineage regulatory factors has been observed and proposed as important for stabilizing repression of such factors and subsequently safeguarding successful iPSC reprogramming. The fact that Dnmt3a/Dnmt3b KO mouse fibroblast reprogram with profoundly lower efficiency has boosted the latter conclusion^33^. We also note that the newly identified de novo DNA methyltransferase DNMT3C^34^ is upregulated during iPSC reprogramming (Extended Data Fig. 8a,b) and thus might be possibly compensating for Dnmt3a/b absence and explain residual iPSC reprogramming in Dnmt3a/Dnmt3b KO MEFs^33^. We characterized DNA methylation dynamics and investigated its connection to histone modifications and transcription regulation. We measured DNA methylation using Whole-Genome-Bisulfite (WGBS) and Reduced-Representation-Bisulfite (RRBS, which is focused on CpG-rich loci) in each time point. We observed a global reduction in DNA methylation (Fig. 3a), which reaches its lowest level at day 8. Intergenic regions, which are typically CpG-poor, are highly methylated in MEF (average methylation of 71%, Extended Data Fig. 8c), and reduced to 42% methylation on day 8. Promoters, on the other hand, are typically CpG-rich, and are lowly methylated throughout the reprogramming. Enhancers show intermediate levels, with average methylation of 54% that is reduced to 13% on day 8.

**Fig. 3:**
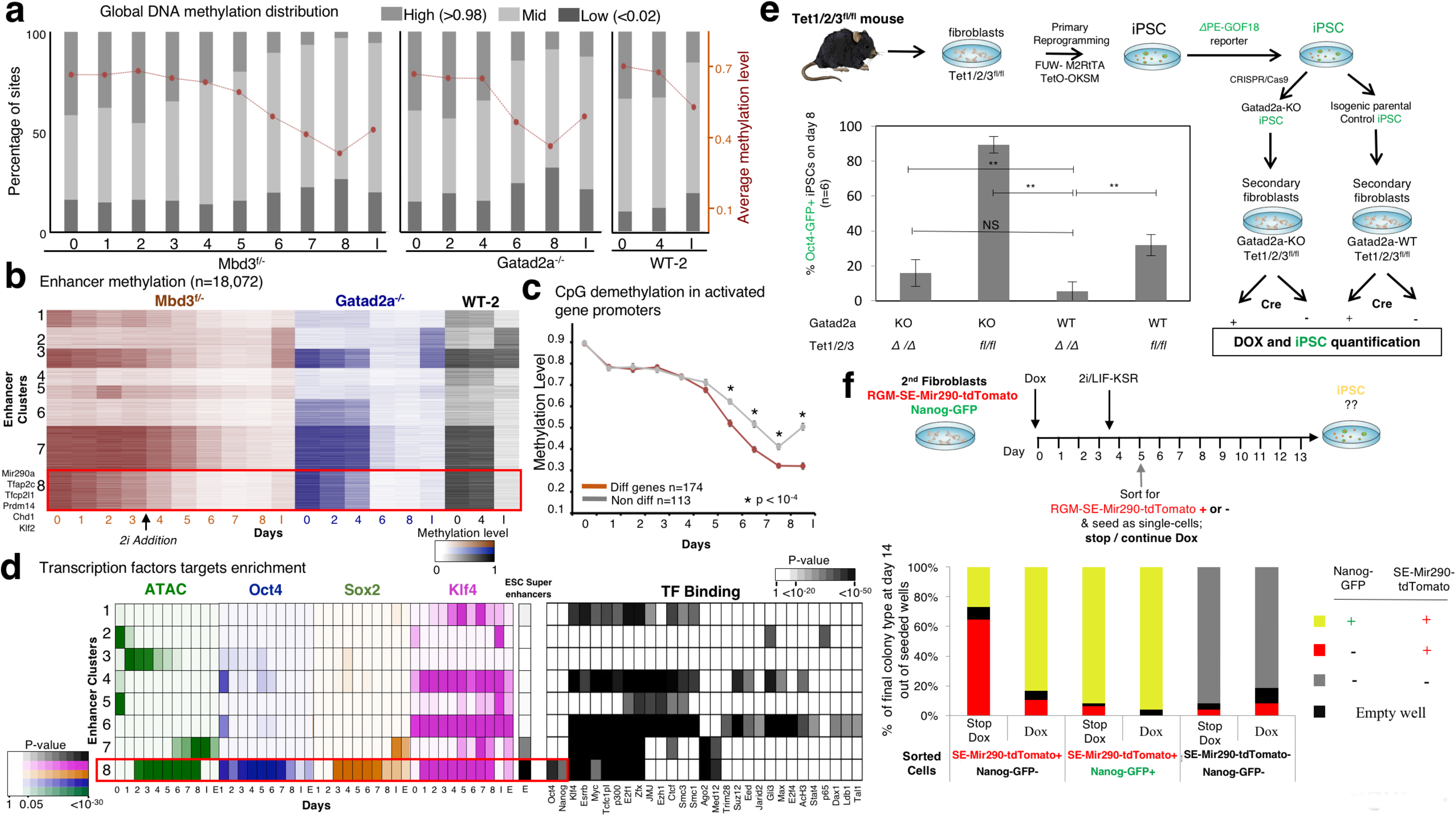
Rapid DNA-demethylation of naïve ESC super-enhancers during conducive iPSC reprogramming. a. Distribution of low (<0.02), mid (0.02-0.98) and high (>0.98) methylated CpG sites, along reprogramming in Mbd3^f/-^, Gatad2a^-/-^, and WT-2 systems. Average and SEM are indicated in red plot. This graph shows a global reduction in DNA methylation that starts early and before 2i introduction in the reprogramming protocol at day 3.5, and reaches its minimal level at day 8 (in Mbd3^f/-^ and Gatad2a^-/-^ systems). b. Methylation level measured in covered enhancers (n=18,072), in Mbd3^f/-^, Gatad2a^-/-^ and WT-2 systems. Enhancers are clustered into eight clusters using k-means. Cluster 8 consists of enhancers that undergo fast demethylation, compared to clusters 3 and 7. c. Average methylation measured in promoters of genes that were highly methylated (>80%) in day0. Genes that change their expression level (red) are compared to genes that do not change their expression level (gray). Wilcoxon p-value indicates places where methylation of differential genes is significantly lower than methylation of non-differential genes. d. Left: Enrichment of enhancer clusters, as shown in panel b, for OSK binding, DNA accessibility, and super enhancers, as defined previously^37^, showing that cluster 8 is highly enriched for OSK binding and overlaps with super enhancers. Color shades represent FDR corrected enrichment p-value. Right: Enrichment of the same enhancer clusters to transcription factor binding, taken from hmChip database^45^. e. Tet1/2/3^fl/fl^ MEFs were reprogrammed to give iPSC, using primary infections of TetO-OKSM lentivirus, followed by Oct4-dPE-Oct4-GFP transgene reporter transduction and validation, and afterwards Gatad2a-KO generation with CRISPR/Cas9. Secondary MEFs were derived from the indicated isogenic lines and subjected to reprogramming. Reprogramming efficiency was measured by Oct4 GFP+ cells percentage in Tet1/2/3 null (Δ) and Tet1/2/3^fl/fl^ with and without Gatad2a expression, after 8 days of reprogramming. Tet1/2/3 null cells reprogramming efficiency was significantly decreased comparing to Tet1/2/3^fl/fl^ cells, in both platforms (Gatad2a-KO and Gatad2a-WT). **p<0.01, ***p<0.001 (Student’s t-test), n=6, error bars indicate SD f. Secondary MEF harboring Mir290-RGM (Reporter of Genomic Methylation) and Nanog GFP-reporter were sorted after reprogramming to 3 different populations: RGM-SE-Mir290-tdTomato positive cells (sorted at day 5), Nanog-GFP and Mir290-RGM positive cells (sorted at days 10-14), and “double negative” cells (sorted at day 5). The cells were seeded as single cell-per-well, and were treated with medium either supplemented with Dox or lacking Dox. On day 14 colonies were inspected for GFP and mCherry (RGM) markers. Above 80% of mir-290-RGM+ cells Nanog-GFP-cells sorted at day 5 were successfully reprogrammed contingent that DOX was continued (i.e. were Nanog-GFP+ on day14).

Next, we clustered enhancers based on their methylation levels in both Mbd3^f/-^ and Gatad2a^-/-^ systems, and in WT-2 (Fig. 3b). All 8 clusters showed different variations of progression exclusively entailing loss of DNA methylation, and none of the clusters showed continuous increase in methylation levels during the 8 days of reprogramming (Fig. 3b). Similar results were obtained for promoters (**data not shown**). The latter indicate that *de novo* DNA methylation is neither required for highly efficient and conducive iPSCs reprogramming and nor for repression of somatic lineage genes in naïve ground state pluripotency reprogramming conditions. This indicates that previously described multiple waves of demethylation and remethylation are not inherent to the success of iPSC process^4,5,9,35^. We wanted to test whether different rates of demethylation exist for certain gene groups. We considered genes that are methylated in MEF (>80% methylation), and compared those that change their expression (FC>4) to those that do not change their expression (FC<1.5, Fig. 3c).

We found that genes which are upregulated at some point during the process, undergo significantly faster demethylation compared to the non-changing genes, starting from day 6 following OSKM induction. Interestingly, when examining the methylation of enhancers (Fig 3b) we identified one cluster (number 8), which is 68% methylated in MEF, and then undergoes fast demethylation (with average 43% methylation level on day 3), even before introducing 2i at day 3.5. The enhancers in this cluster are accessible between days 2 and 8, are highly enriched for the binding of OSK (Fig. 3d), and highly overlap (p<10^-43^) with ESC super-enhancers^36^ including Mir290a, Tfcp2l1, Tfap2c, Bend3, Klf2, Prdm14, Tet3, Chd1, n-Myc and Foxo1, which are known to boost reprogramming efficiency (**Supplementary Table 5**)^37^. Another cluster of enhancers (cluster 7) is enriched for SK binding and for ESC super enhancers, but it undergoes a slightly slower demethylation than cluster #8 (with average methylation of 67% in day 3), and its enhancers are accessible as measured by ATAC-seq only from day 6 until iPS/ES (Fig. 3d). Both clusters are enriched for binding by Esrrb, E2f1, Klf4 (Fig. 3e), but only cluster number 8 is enriched for Nanog and Oct4 binding.

We next aimed to unravel the mechanism underlying these different demethylation rates in our system and whether this early demethylation is important for achieving highly efficient reprogramming upon optimized NuRD depletion. Given that this demethylation occurs before introducing 2i we suspected that Tet enzymes, known to target and demethylate key pluripotency genes in ESC^38^, might regulate this change. To test this, we established Tet1/Tet2/Tet3 triple floxed conditional knockout mouse model (Extended Data Figure 9a-e), from which we derived secondary iPSCs, generated isogenic Gatad2a^-/-^ iPSC lines with CRISPR/Cas9 and subsequently re-isolated DOX inducible reprogrammable MEFs (Fig. 3e). Depletion of Tet enzymes in Gatad2a-WT decreased iPSC efficiency (from 32% to 6% efficiency), consistently with previous reports showing that overall Tet enzyme are dispensable for iPSC formation^39^. However, upon ablation of Tet enzymes in the Gatad2a^-/-^ deterministic reprogramming system, reprogramming efficiency dropped from 93% down to 6-18%, similar to that in WT system, thus abolishing the beneficial effect of Gatad2a depletion (Fig. 3e). The latter indicate that Tet activity early in reprogramming is essential for highly efficient conducive iPSC reprogramming in NuRD depleted systems. To test whether early demethylated enhancers of genes in cluster 7 and 8 functionally contribute to conducive reprogramming, siRNA depletion of selected members of cluster 8 genes, including Tfap2c and Tfcp2l1, was applied. Knockdown of Tfap2c or Tfcp2l1, but not Bend3, reduced reprogramming efficiency by 15% (Extended Data Fig. 9f). The latter suggests that early demethylation of selected enhancers by Tet enzymes promotes the commissioning of several pro-reprogramming factors that synergistically contribute to the highly efficient reprogramming observed herein.

Finally, we tested whether the rapid demethylation of cluster 8 super-enhancers specifically detected during deterministic iPSC reprogramming, but not in measurement conducted on bulk WT reprograming samples (Fig. 3b), can be used as an early marker to prospectively enrich for the rare correctly commissioned WT cells to become iPSCs. To isolate cells in real time during reprogramming based on their DNA methylation status of a certain locus and at the single cell level, we utilized a recently generated reporter system for endogenous genomic DNA methylation (RGM)^40^. We chose a validated RGM construct for Mir290 super-enhancer encoding tdTomato (RGM-SE-miR290-tdTomato), which was enriched in cluster 8, and introduced it in two OKSM DOX inducible secondary reprogramming systems carrying knock-in Nanog-GFP reporter that specifically marks iPSC generation (Fig. 3f). In these relatively inefficient Gatad2a and Mbd3 WT systems, the first Nanog-GFP+ cells appeared at days 10-14 following DOX induction, which were sorted and plated as single cells in naïve ESC media with or without continued DOX. As expected, over 90% iPSC efficiency was obtained following sorting Nanog-GFP+ cells irrespective to the continued use of DOX to induce transgenes after sorting (Fig. 3f), confirming that Nanog-GFP+ cells are already bona fide committed iPSCs that no longer need OSKM transgene expression. On the contrary, SE-miR290-tdTomato+ cells appeared at very low frequency already at day 4-5 during reprogramming of Mbd3/Gatad2a-WT cells as single positive tdTomato+ cells (Extended Data Fig. 9g). tdTomato+/GFP-cells at day 5 were sorted and plated as single cells in naïve ESC media with or without continued DOX treatment. Remarkably, >85% iPSC efficacy, as measured by Nanog-GFP, was obtained from day 5 sorted tdTomato+/GFP-cells only upon continued DOX supplementation (Fig. 3f, Extended Data Fig. 9h). In the absence of continued DOX, 26% efficiency was obtained from early tdTomato+/GFP-sorted cells (Fig. 3f), suggesting that the sorted tdTomato+/GFP-cells are not bona fide iPSCs, however they were correctly “commissioned” and become committed to becoming iPSCs if OSKM expression is continuously delivered to drive the process toward completion. Day 5 double negative sorted cells did not yield any iPSCs after 10 days of DOX induction (Fig. 3f), indicating that this fraction marks somatic cells that did not optimally embark on a conducive trajectory towards becoming iPSCs. These results indicate that early demethylation of Mir290 super-enhancer marks correctly commissioned NuRD-WT somatic cells following DOX induction, that rapidly assume a conducive trajectory to becoming iPSCs if OKSM induction is continued. This also provides a means to prospectively isolate adequately commissioned somatic cells for a successful reprogramming trajectory based on endogenous epigenetic feature, rather than cell surface markers which relatively have much lower predictive fidelity.

### Two synergistic and distinctly regulated gene programs ignite conducive reprogramming

We next wanted to characterize the epigenetic changes and examine their connection to the changes in gene expression. For each differentially expressed gene that showed a significant epigenetic modification in its promoter (n=7,801), we calculated the correlation between its transcriptional temporal pattern and chromatin modification patterns, measured around the transcription start site or transcription end site (TSS and TES, respectively). When we cluster these genes and chromatin marks (Fig. 4a), we observed that chromatin marks separate into two clusters: One consists of marks which are positively correlated to gene expression, and are indeed known to be associated with active transcription, such as H3K4me3^41^, H32K7Ac^42^, H3K36me3^43^ (in TES) and chromatin accessibility (measured by ATAC-seq). The other consists of marks which negatively correlate with gene expression, and are known repression-associated marks, such as H3K27me3 and H3K9me3^44^. Interestingly, the genes also separate into two main clusters. One consists of genes that display high correlation (positive or negative) between expression and chromatin modifications, and the other consists of genes that are not correlated, despite the fact that the genes are differentially expressed. Notably, each of these two gene groups contains both induced and repressed genes (Extended Data Fig. 10a,b). We inspected the actual transcriptional and epigenetic patterns for these two gene clusters, focusing on H3K27ac and H3K27me3 marks, which showed the highest positive and negative correlation to transcription (Fig. 4b). The genes in the first group showed a clear switch-like behavior between the epigenetic marks (**upper panels in** Fig. 4b, Extended Data Fig. 10a), correlated with the activation or repression. We therefore concluded that these are genes with Epigenetically Switched Promoters (abbreviated as ESPGs). In the second group, the majority of the genes (N=3049, 72%) had differential transcription (above 4-fold change), but with consistently high levels of H3K27ac and low levels of H3K27me3 (z-score <0.7, **lower panels in** Fig. 4b, Extended Data Fig. 10b). The promoters of these genes show a constitutive active chromatin signature, suggesting that these genes are regulated by distinct mechanisms. We refer to this group as CAPGs (Constitutively Active Promoter Genes) (**Supplementary Table 6**). In accordance with chromatin modifications, DNA methylation in the promoters of the two groups is different (Extended Data Fig. 10c): CAPGs show a consistent hypomethylation, regardless of their transcriptional pattern, whereas ESPGs, which are regulated on the chromatin level, are also regulated by DNA methylation (i.e. ESPGs which are repressed in MEF and are upregulated during the process show a decrease in their DNA methylation level– Extended Data Fig. 10c). The same epigenetic phenomenon was observed in the enhancers associated with ESPGs and CAPGs (Extended Data Fig. 11a).

**Fig. 4:**
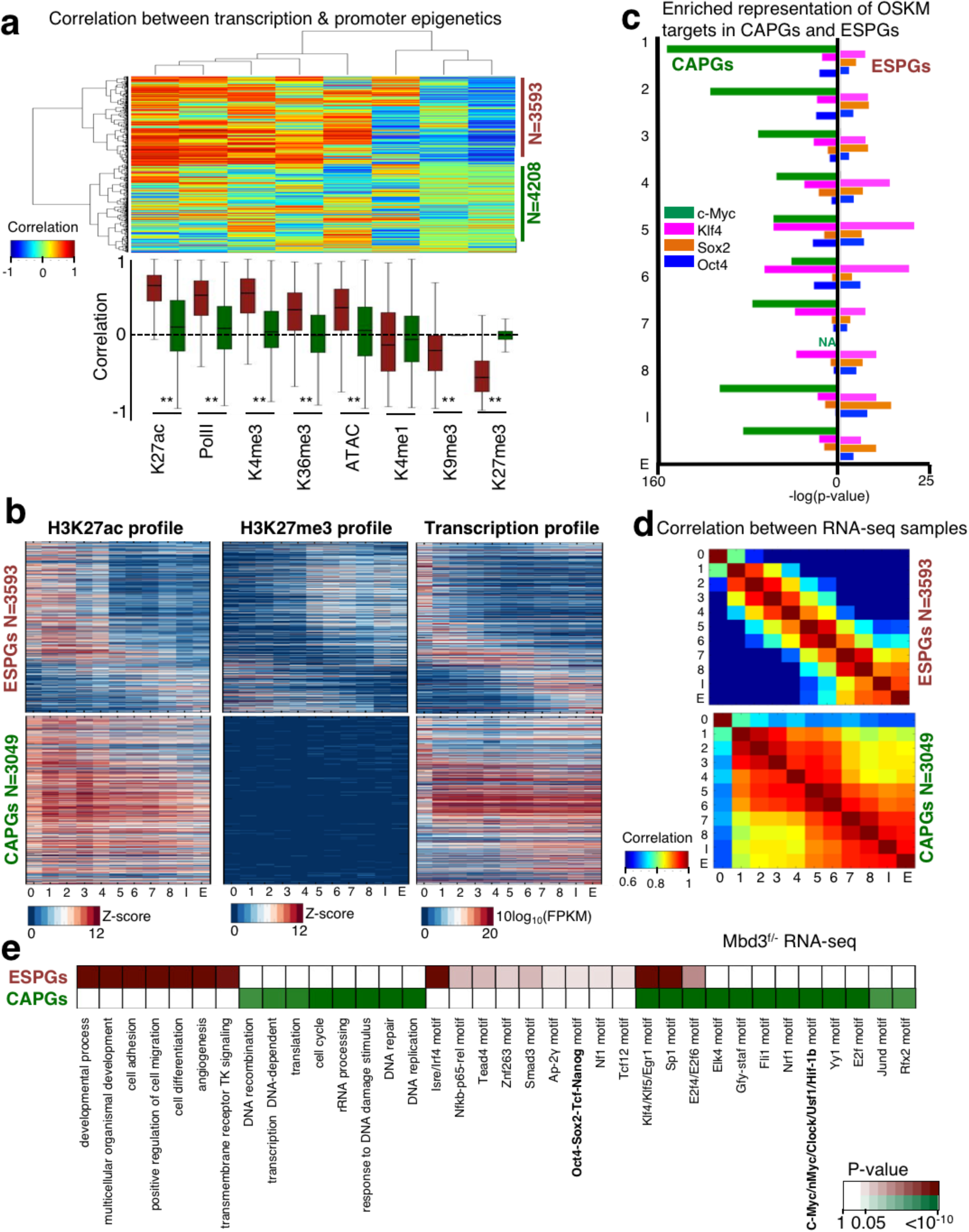
Distinct regulation of cell fate genes and of biosynthetic processes. a. Spearman correlation between the expression change of each differential gene (row) and change in promoter chromatin modification of each indicated mark (column). Analysis was done on all differential genes that have at least two marks in their promoter (i.e. with z-score>1 std), resulting in 7801 genes. Hierarchical clustering clustered the genes into two distinct groups: genes with correlation between gene expression and promoter epigenetic modifications (n=3593) and genes with no trend of correlation (n=4208). Top: clustered correlation matrix. Bottom: correlation distribution of each mark in the two gene groups, where red denote correlated group, and green denote the non-correlated group. ** Wilcoxon p-value < 10^-170^ b. Pattern of H3K27ac, H3K27me3 and expression in Epigenetically Switched Genes (ESPGs, top), and in genes with Consistency-Active-Promoters (abbreviated as CAPGs, bottom). Each row corresponds to a single gene, genes are sorted according to their expression pattern and the same sorting was applied to the epigenetic marks. c. Enrichment of OSKM binding targets in promoters of ESPGs compared to CAPGs. Minus log_10_ of Fisher exact test p-values are indicated. d. Spearman correlation matrix between Mbd3^f/-^ RNA-seq samples calculated over ESPGs (Top), showing a gradual change along reprogramming and over CAPGs (Bottom), showing two waves of change, on day 1, and on days 5-6. e. Enrichment of GO categories and binding motifs in ESPGs (red), CAPGs (n=3049, green). Color shades indicate FDR corrected Fisher exact test p-value, or motif enrichment p-value.

Inspecting the functional enrichment of the two groups, we found a specific association of ESPGs to cell fate determination processes, indicating that epigenetic regulation is highly specific for cell fate genes. CAPGs are enriched for biosynthetic pathways including DNA synthesis, proliferation, DNA repair and chromatin reorganization (Fig. 4e, Extended Data Fig. 11b). The two programs show a distinct conservation pattern during the evolution of vertebrate organisms: while in vertebrates CAPGs and ESPGs are conserved in a similar degree (Extended Data Fig. 12a), in fungi and other non-vertebrates CAPGs are more conserved than ESPGs (p<10^-30^), emphasizing their basic role in cellular maintenance. The two groups also show distinct regulation by c-Myc: CAPGs, but not ESPGs, are significantly bound by c-Myc (sample median p<10^-75^ for CAPGs, sample median p>0.9 for ESPGs) (Fig. 4c). This is supported also by over representation of c-Myc motif only in CAPGs promoters (Fig. 4e). Additional TF binding motifs show enrichment specific to one group and not the other (Fig. 4e), further supporting a model of separate regulation. Finally, the two groups have different temporal behavior: while ESPGs have a gradual change in activity along reprogramming, CAPGs converge to their final activity pattern as early as day 1 (Fig. 4b, d). Importantly however, these two programs retain a coupled and cross-coordinated regulation. Protein binding enrichment in ESPGs and CAPGs using public protein-DNA databases^45-47^ shows a number of proteins that are associated with one of the groups, but bind the opposite group (Extended Data Fig. 12b). For example, several epigenetic modifying components such as Polycomb (Suz12, Eed, Ezh2), Wdr5, methylation associated family (Brca, Ddb2, Tet1, Tet2), Sirt family (Sirt1, Sirt3) and Zfp281, all showing a constitutively active promoter configuration, but regulate ESPGs (Extended Data Fig. 12b).

### Two divergent modes of epigenetic repression of ESPGs during conducive reprogramming

We next sought to discern epigenetic regulation during iPSC formation, and from the differentially expressed genes we focused on ESPGs as they are the ones that undergo a repressive to activation switch or vice versa. We used the chromatin modification coverage in promoters and the expression level we obtained with RNA-seq, and calculated the temporal correlation distribution for all ESPGs (Fig. 5a), i.e. correlation that is calculated for each gene, over all time points. RNA-PolII, H3K27ac and H3K4me3 in promoters are highly correlated to gene expression of the genes they decorate (Median r=0.55,0.7,0.6, respectively). H3K27me3, H3K9me2 and H3K9me3 show negative correlation to gene expression. H3K27me3 also shows high anti-correlation to H3K27ac, since these are two mutually exclusive marks occurring on the same lysine residue^42,48^. Examining the frequencies of combinations of chromatin modifications (Fig. 5b) on gene promoters, we observed that in upregulated ESPGs (n=431), there is a rapid reduction of H3K9me2 and H3K27me3. In addition, there is a substantial increase in H3K27ac and binding of PolII, such that by day 8 and iPSC, 45% of the promoters are decorated by the combination of H3K27ac and PolII. In the downregulated ESPGs (n=974), we observe the opposite pattern with loss of H3K27ac and PolII binding, and gain of H3K27me3 starting from day5 (See green and red parts in Fig. 5b **– right panel**).

**Fig. 5:**
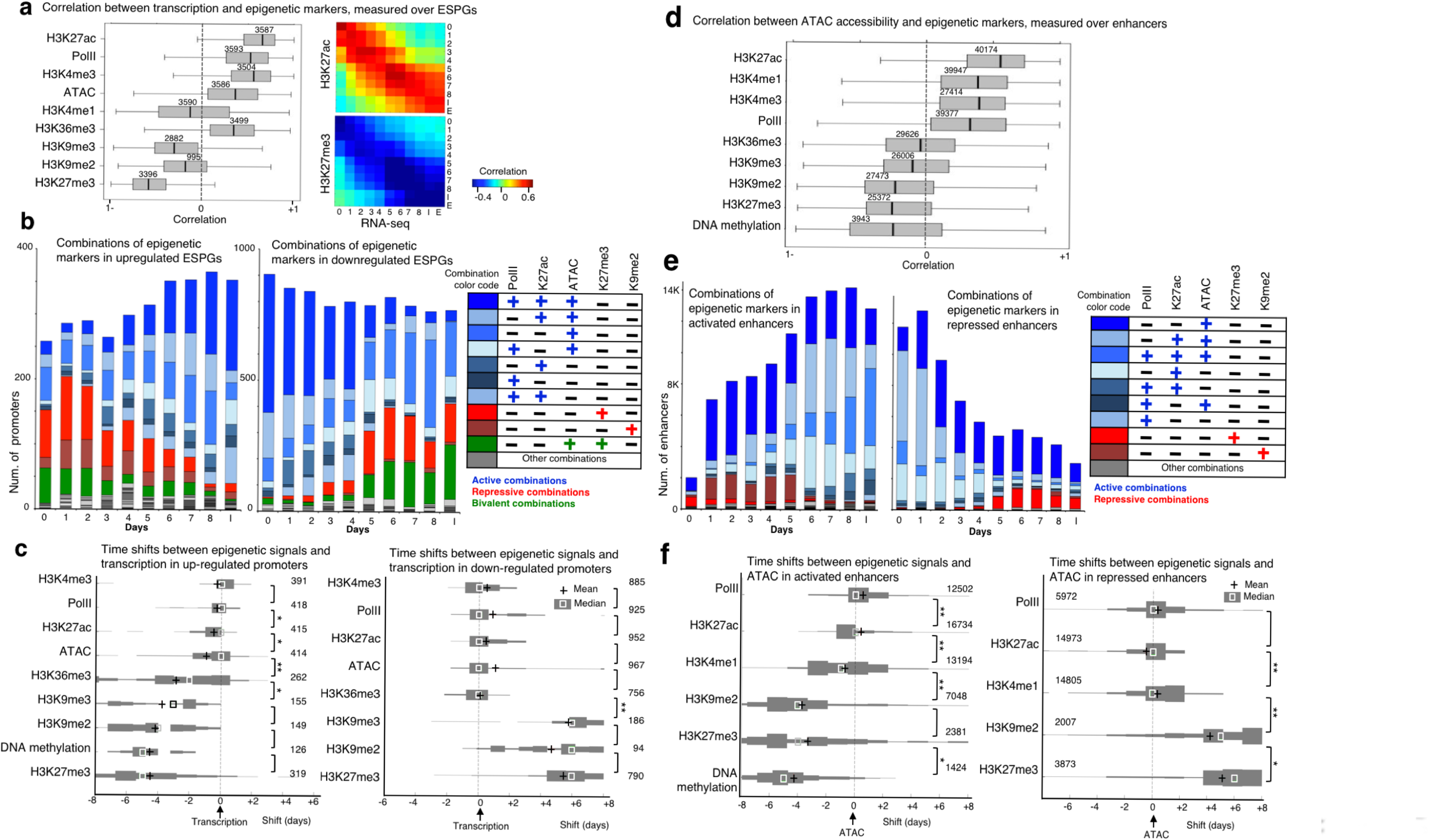
Mapping the order of epigenetic events that drive transcription initiation and repression. a. Left: Correlation between each of the indicated chromatin modifications and gene expression patterns that were measured in the promoters of ESPGs that have the modifications (numbers are indicated). Correlations were calculated for each gene, over 11 time points. H3K27ac, PolII and H3K4me3 show positive correlation with gene expression, while H3K27me3 and to lesser extent H3K9me2,3 show negative correlation with expression. Right: Correlation matrix over all ESPGs, between H3K27ac or H3K27me3 and gene expression (RPKM) for each sample separately. b. Stacked bar chart of all combinations of the indicated chromatin modifications, as measured in promoters of upregulated ESPGs (left) and downregulated ESPGs (right). Right– color code of frequent combinations (>3% of each sample). Note that H3K27ac and H3K27me3 are mutually exclusive (frequency<2%), and so are H3K27me3 and H3K9me2. Upregulated genes move from repressive combinations (K27me3/K9me2), to active combinations (PolII/K27ac/ATAC) while downregulated genes gain repressive combinations. c. Distribution of time shifts between each of the indicated epigenetic modification and gene expression profile, measured using cross correlation. The distribution, presented as histogram, was measured over upregulated ESPGs (Left) and downregulated ESPGs (right) which have a changing epigenetic modification (max-min z-score > 0.5). The number of promoters tested is indicated. Plus indicates mean, and square indicates median. *p<10^-5^, **p<10^-25^ (Wilcoxon test). d. Correlation between each of the indicated chromatin modification and DNA accessibility patterns that were measured in all differential enhancers that have the modifications (numbers are indicated). Correlations were calculated over 11 time points. H3K27ac, H3K4me1, H3K4me3 and PolII show positive correlation with DNA accessibility, while DNA methylation, H3K27me3 and H3K9me2 show negative correlation with accessibility. e. Stacked bar chart of all combinations of the indicated chromatin modifications, as measured in activated enhancers (left) and in repressed enhancers (right). Right– color code of frequent combinations (>3% of each sample). Note that H3K27me3 and H3K9me2 are mutually exclusive. Activated enhancers move from repressive combinations (K27me3/K9me2 or no mark), to active combinations (PolII/K27ac/ATAC) while repressed enhancers undergo the opposite process. f. Distribution of time shifts between each of the indicated epigenetic modification and accessibility profile, measured using cross correlation. The distribution, presented as histogram, was measured over activated enhancers (Left) and repressed enhancers (right) which have a changing epigenetic modification (max-min z-score > 0.5). The number of enhancers tested is indicated. Plus indicates mean, and square indicates median. *p<10^-4^, **p<10^-50^ (Wilcoxon test).

The analysis above also highlights the most frequent combinations, and the combinations that are not apparent in the data and are mutually exclusive. The latter allowed us to ask whether mutually exclusive modes of repression exist in iPSC reprogramming. Active marks (H3K27ac, RNA-PolII, ATAC) tend to appear together on promoters (Fig. 5b), and we did not discern distinct mutually-exclusive modes of acquiring activation marks. On the contrary, repressive marks (H3K27me3, H3K9me2) work separately from one another. We observed that less than 1% of the promoters are marked by both H3K9me2 and H3K27me3 (Fig. 5b), suggesting these are mutually exclusive marks. Indeed, our data show a clear association between H3K9me2 and DNA methylation (Extended Data Fig. 13a) and this may explain why in our system, which undergoes substantial DNA demethylation (Fig. 3a), there is very limited gain of H3K9me2 on downregulated ESPGs. Furthermore, H3K27me3 decorates genes that are enriched for functions in development, while H3K9me2 decorates genes related to signaling pathways (Extended Data Fig. 13b). H3K27me3 genes are naturally highly enriched for Polycomb targets^49^ (Extended Data Fig. 13b), and an induction in the expression of Polycomb members is observed, which overlaps with the increase in H3K27me3 peaks starting from day 5 (Fig. 5b, Extended Data Fig. 3b). Altogether, this analysis uncovers two divergent modes of epigenetic repression by H3K27me3 and H3K9me2 during iPSC reprogramming with opposing association with DNA methylation and distinct associated regulatory functions.

Bivalent promoters, which carry both H3K27me3 repressive mark and H3K4me3 active mark, constituted 38% of ESPGs and were found to constitute a third distinct mode of repression (green combination in Figure 5b and Extended Data Fig.13f). Bivalent promoters are highly enriched for developmental regulators (Fisher exact test p-val<10^-90^, FDR corrected), and overlap with bivalent promoters that were detected previously in MEFs and in ESCs^4^ (Extended Data Fig. 13b). When comparing bivalent promoters to H3K27me3-only promoters we observe that the repression of transcription is stronger in H3K27me3-only promoters than in bivalent ones (One tailed Wilcoxon test, p-value<10^-98^). Moreover, the chromatin of bivalent promoters is much more accessible compared to H3K27me3-only or to H3K9me2 promoters, which decorate closed chromatin (Extended Data Fig. 13c,d,f). To rule out the possibility that bivalent signature is a mere result of a residual mixed cell population in the highly efficient system, we note that other combinations that are mutually exclusive in promoters, such as H3K27me3 and H3K27ac, appear in much lower frequency (<3%) compared to the bivalent combinations (>25%). The latter is further supported by the observed difference between promoter and enhancer bivalent probability (Extended Data Fig. 13e,f).

### Repressive and active chromatin mark switching is temporally separated over ESPGs

The temporal interplay between the different chromatin marks remains to be defined at high-resolution, and it is unclear whether during transitions from repressed to activated state (or vice versa) changes in repressive and activating epigenetic modes co-occur simultaneously or are well separated. Since our data consist of time-series we used Cross Correlation, a signal-processing algorithm widely used to detect and quantify the temporal offset between signals^50,51^ (Extended Data Fig. 14), to test whether the deposition of these modifications has a temporal order. We estimated the distribution of offsets across ESPGs (Fig. 5c), i.e., for each ESPG we calculate the cross-correlation between its temporal expression and the temporal pattern of each of the epigenetic marks. The analysis clearly highlighted separation between accumulation of repressive and activation marks at gene promoters. In induced genes, first, DNA is demethylated and H3K27me3 is removed, and only then chromatin becomes accessible. Finally, H3K27ac, RNA-PolII and H3K4me3 accumulate in the promoter in a close proximity to transcriptional activation (Fig. 5c**, left**). The latter also excludes an alternative scenario wherein gene activation, removal of repressive marks follows epigenetic activation and transcription initiation (Fig. 5c). In repressed genes, PollI disassociates from its bound promoters in close proximity to the eviction of H3K27ac and H3K4me3 and chromatin closure (Fig. 5c**, right**). Only afterwards, repressive marks like H3K27me3 and H3K9me3 gradually accumulate during the following days.

Our next aim was to elucidate the temporal order of epigenetic changes that occur in differential enhancers (Extended Data Fig. 4) and how they compare to those observed in promoters. 43% of all annotated enhancers (n=40,174) showed differential ATAC-seq and H3K27ac signals in both Mbd3^f/-^ and Gatad2a^-/-^ systems, and were identified as differential enhancers (Extended Data Fig. 4b). The enhancer activation kinetics in the two NuRD depleted systems were highly consistent and faster than the WT systems (Extended Data Fig. 5a-c). We systematically calculated the correlation between chromatin accessibility and chromatin modification in each of the differential enhancers (Fig. 5d), and observed positive correlation with H3K27ac modification (median r=0.55), and unlike in promoters, positive correlation with H3K4me1 modification (median r=0.4). Interestingly, positive correlation was evident between enhancer accessibility and RNA-PolII binding. Furthermore, we observed negative correlation between enhancer accessibility and DNA-methylation, H3K27me3 and H3K9me2, but to less extent with H3K9me3 modification. Interestingly, H3K27me3 does not always decorate repressed enhancers. In fact, when all possible combinations of chromatin marks are inspected in differential enhancers (Fig. 5e), 85% of the enhancers which are active in day 8 are in a closed chromatin state on day 0 (MEF), but are not marked by any of the histone marks measured herein. Like in promoters, H3K9me2 repression can be observed in the first days of reprogramming, is later depleted, and is mutually exclusive to H3K27me3 (Fig. 5e). Unlike its abundance on promoters during reprogramming, bivalency at enhancers (H3K27me3 with H3K4me1) is rare, and H3K27me3 is rarely deposited on accessible enhancers (**<4%**, Fig. 5e; Extended Data Fig. 13e).

To examine the sequence of epigenetic events during enhancer activation and suppression, we again used Cross-Correlation and quantified the temporal offset between chromatin changes and DNA accessibility in each differential enhancer. We found that in activated enhancers (n=17,174), H3K27me3 is first removed, then H3K4me1 is deposited, followed by chromatin accessibility and deposition of H3K27ac and finally, by binding of RNA-PolII (Fig. 5f). In repressed enhancers, PolII release and the removal of H3K27ac and H3K4me1 happen all in close proximity to chromatin closure, followed by gradual deposition of H3K27me3 or H3K9me2. Thus overall, the orderly switches from activation to repression (or vice versa) over enhancers are similar to those seen over promoters (Fig 5c,f).

Since enhancers control transcription, one may expect to see a connection between the epigenetic state of a certain enhancer and the promoter it regulates. We then used Cross-Correlation to quantify the temporal order of epigenetic changes in enhancers and promoters in relation to measured transcription changes (Extended Data Fig. 15a). No significant temporal differences were observed in deposition or removal of repressive chromatin marks between enhancers and promoters during repression or activation of ESPGs, respectively (Extended Data Fig. 15a). However, we could see that active modifications are deposited on enhancers before they are deposited on the associated promoters during gene activation (paired sample t-test ATAC-seq p<10^-7^, H3K27ac p<10^-11^, H3K4me3p<10^-2^, respectively). In contrast, during ESPG repression, eviction of activation marks on enhancers was significantly lagging in comparison to promoters (Extended Data Fig. 15a). Unexpectedly, RNA-PolII binds enhancers and showed similar behavior to the activating epigenetic marks (PolII binds enhancers slightly before it binds to promoters (p<10^-3^, Extended Data Fig. 15a,c) and leaves the enhancers slightly after it leaves the promoters (p<10^-23^))^52^. RNA-PolII binding in enhancers is highly correlated both to gene transcription (Extended Data Fig. 15d) and to enhancer activity (Fig. 5d). Independent RNA-PolII binding data, measured in mouse ESC^27^, was also highly enriched among enhancers which are active in late reprogramming stage (p=<10^-200^, Extended Data Fig. 15b), and the same is true for binding of Nelfa, a member of RNA-PolII elongation complex. These results indicate that the phenomenon of PolII recruitment to enhancers as an early event of enhancer commissioning, is widely abundant during iPSC reprogramming. Whether and how PolII is functionally involved in enhancer-gene interaction is of future scientific interest.

### Myc activity is essential for iPSC reprogramming

While the above sections focused on dissecting functional and molecular dynamics of ESPGs, we now turned to assess the regulation and importance of CAPG changes in facilitating successful reprogramming. As indicated earlier (Fig. 4c, e), CAPGs are predominantly regulated by Myc and drive cellular biosynthetic processes. As exogenous Myc is dispensable for iPSC formation from WT and NuRD-depleted somatic cells^53,54^, this raised the possibility that the observed CAPGs induction is merely a side-effect of c-Myc over-expression and is not essential for the reprogramming process. To tackle this question, we introduced a number of perturbations to the highly efficient optimally NuRD-depleted reprogramming protocols. First, we tested reprogramming with a viral induction of only 3 factors OSK (without c-Myc) (Fig. 6a (**i**)). Notably, CAPGs that were upregulated in the original protocol, were still significantly upregulated compared to MEF (P-val<10^-12^ Extended Data Fig. 16a). However, we noticed that in OSK reprogramming, endogenous c-Myc continues to be highly expressed and endogenous n-Myc is induced after OSK induction (FC>1.8, for both c-Myc and n-Myc). We therefore tested OSK reprogramming under inhibition of endogenous Myc family members by treating MEFs that carry OSK cassette with siRNAs for c-Myc, n-Myc and l-Myc starting on day −3 prior to DOX induction (Fig. 6a (**ii**)). Myc inhibition resulted in dramatic reduction in reprogrammed colonies (Fig. 6b,c) and greatly reduced induction and repression of CAPGs (Fig. 6c). Surprisingly, the downregulation and upregulation of ESPGs was also diminished by Myc inhibition (Fig. 6c), although Myc does not bind them directly (Fig. 4c,e); suggesting that this change is likely caused by an indirect effect. Inhibition of Myc activity with pharmacological inhibitor 10058-F4^55^ yielded a similar result to siRNA experiments (Fig. 6a (**iii**),d and Extended Data Fig. 16b,c)). Finally, we used conditional knockout fibroblasts for both c-Myc and n-Myc genes and carrying Lox-stop-Lox-YFP reporter in the Rosa26 locus which can mark floxed cells upon Cre-treatment. Fibroblasts were treated with CAGGS-Cre plasmid, sorted for YFP and subjected to either OSK or OSKM transduction (Fig. 6e). Remarkably, we could not obtain any YFP+ iPSC colonies following OSK induction and follow up of over 30 days of reprogramming from Cre–treated cells (Fig. 6f). Isogenic control cells that were not treated with Cre expectedly yielded iPSC following OSK transduction (Fig. 6e,f). OSKM transgenes yielded iPSCs irrespective to the endogenous Myc genotype, consistent with c-Myc transgene ability to compensate for endogenous lack of Myc (Fig. 6f).

**Fig. 6:**
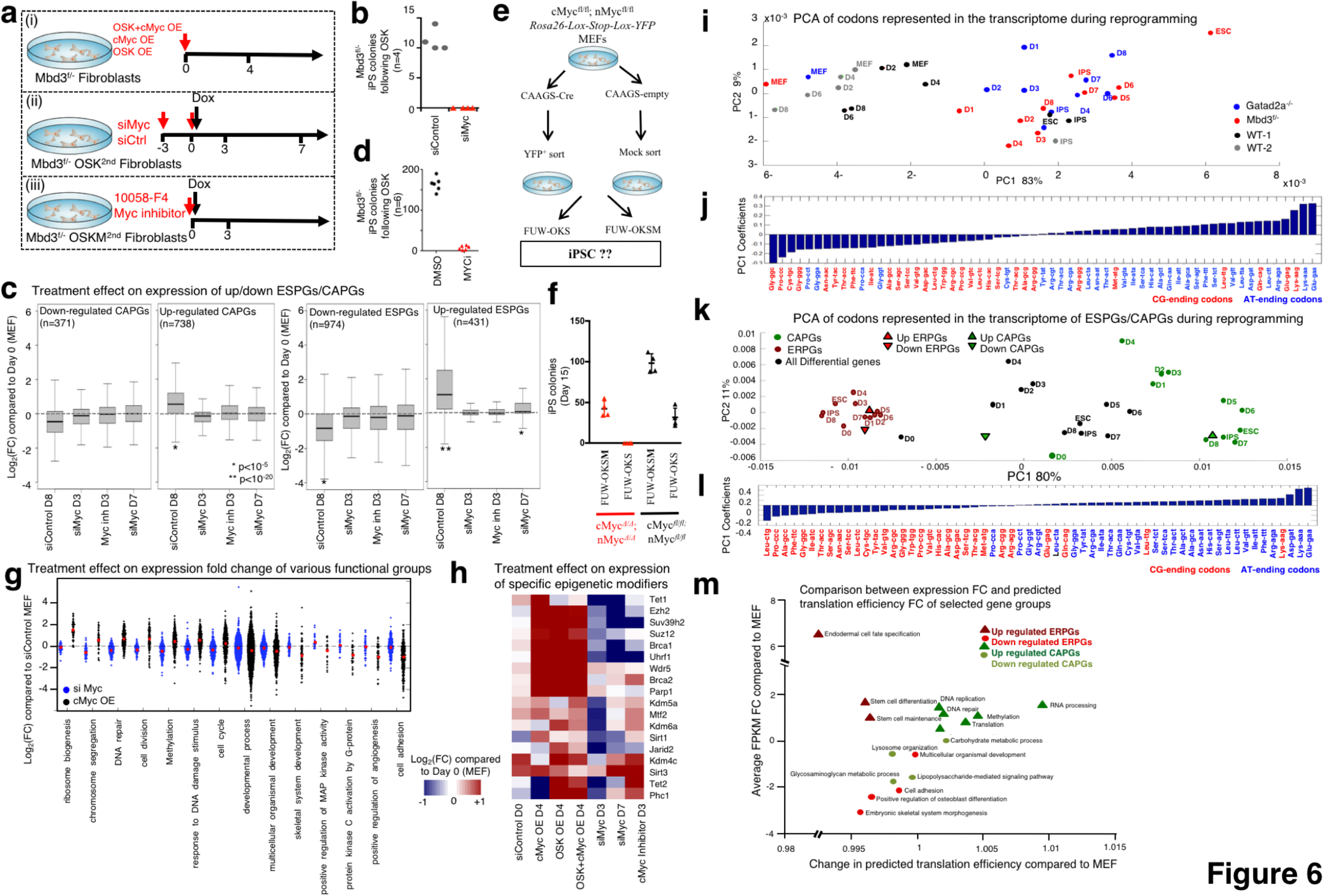
Biosynthetic processes are regulated by endogenous Myc activity and coordinated optimization of available tRNA pool. a. Experimental flow describing three experimental perturbation settings: (i) Mbd3^f/-^ MEFs were virally infected with cMyc over-expression (OE) cassette, OSK-OE cassette or both cassettes. Gene expression was measured on day4 following infection. (ii) Mbd3^f/-^ MEFs carrying OSK Dox-dependent cassette were treated for knockdown of c-Myc, n-Myc and l-Myc. Gene expression was measured on days 3 and 7, and colony formation was measured on day 11. (iii) Mbd3^f/-^ MEFs carrying OSKM Dox-dependent cassette were treated with inhibitor of cMyc (10058-F4^55^) and with Dox. Gene expression and colony formation were measured on day 3. b. Reprogrammed colony formation in Myc knockdown, measured 11 days after Dox induction. c. Distribution of expression fold change (FC, in log_2_ scale) compared to MEF of up/down regulated ESPGs and CAPGs. Presented perturbations are Myc knockdown, inhibition of Myc activity with small molecular inhibitor (10058-F4). (*p<10^-5^, **p<10^-20^, Wilcoxon test) d. Reprogramming efficiency following MYC-inhibitor treatment, measured 12 days after Dox induction. e. Experimental scheme describing reprogramming experiments which were conducted on the background of cMyc and nMyc conditional knockout. f. iPSC Reprogramming efficiency in different cells expressing both endogenous and/or exogenous cMyc and nMyc g. Expression fold-change distribution (log_2_ scale) of selected GO categories in Myc over-expression or Myc knockdown, showing that upon over-expression of Myc, processes such as ribosomal biogenesis and chromosome segregation are induced. h. Expression fold change in the indicated conditions compared to MEF, of selected chromatin modifiers, showing the induction of some of them by the mere over-expression of Myc. i. A PCA projection of codons’ representation in the transcriptome along reprogramming in Mbd3^f/^, Gatad2a^-/-^ and the two WT systems. The representation of the codons in the transcriptome was determined by multiplying the number of occurrences of each codon in each gene by the scaled expression level of each gene in each time point/cell type. The variance percentage, out of the total original variance in the high-dimensional space, spanned by the first and second PCs is indicated on the x and y axis, respectively. j. Coefficients associated with first principle component (Fig 6i, x-axis). Blue– A/T ending codons, Red– G/C ending codons. k. A PCA projection of the codon usage of all differential genes (black), ESPGs (red) and CAPGs (green) show a striking separation between ESPGs and CAPGs. l. Coefficients associated with first principle component (panel a, x-axis). m. Comparison between predicted changes in translation and changes in transcription for various GO categories that show the highest association with up/downregulated CAPGs/ESPGs. CAPGs and downregulated ESPGs show correlated changes in transcription and translation, where upregulate ESPGs show increased expression and predicted reduction in translation efficiency.

Molecularly, c-Myc over-expression (OE) in MEF, without the induction of other reprogramming factors, induced CAPGs expression changes in the same way it changes during reprogramming by OSKM (Extended Data Fig. 16a), also causing significant repression of downregulated ESPGs (somatic genes), but did not lead to the induction of upregulated ESPGs (pluripotency genes). We further validated Myc induced CAPG changes by looking at specific functional groups of genes: Genes related to cell biosynthesis (e.g. ribosomal genes, cell cycle), which are directly bound by c-Myc (Fig. 6g, Extended Data Fig. 16d) are induced upon overexpression of c-Myc. These expression changes are consistent with previously published data^55^ of Myc inhibition and reconstitution measured independently during naïve mouse ESCs maintenance (Extended Data Fig. 16e). Interestingly, we observed that reprogramming related chromatin modifiers such as Prc2 members (Suz12, Ezh2), Tet1, Wdr5 are induced by the mere OE of c-Myc, and fail to be induced upon its inhibition (Fig. 6h). This indicates that Myc has a critical role far beyond the previously described function in repressing somatic genes within the ESPGs^25^. This pertains to igniting the biosynthetic pathways that are dispensable for pluripotency maintenance^55^, yet essential for reestablishing pluripotency in somatic cells and must be provided either endogenously and/or exogenously (in both NuRD WT or depleted systems).

### Rapid rewiring of tRNA pool boosts CAPG

The above results elucidated promotion of CAPG transcription by direct activation by Myc. However, the rapid change in CAPGs expression, without associated changes in their epigenetic signature, raised the possibility that CAPGs may be regulated post-transcriptionally. A recent study^56^ documented a cancer formation promoting mechanism that supports loss of somatic identity and acquisition of a highly active metabolic state during cancer transformation involving coordinated changes in the tRNA pool and the codon usage preference of tRNA. We thus examined if such shifts occur at the codon usage level of the transcriptome and at tRNA transcription status when somatic cells undergo reprogramming toward pluripotency. To characterize putative changes in the codon usage of the transcriptome, we calculated the average codon usage distribution of all differential genes in four reprogramming systems (Mbd3^f/-^, Gatad2a^-/-^, WT-1 and WT-2). Using PCA we characterized the codon combination that shows the highest variability during reprogramming (Fig. 6i), and noticed a change in codon combination that separates between early and late stages of reprogramming. Interestingly, the observed change in codon usage corresponds to a shift from G/C-ending codons (red color) to A/T-ending codons (blue color) (Fig. 6j), with the most prominent change occurring already at the first day of reprogramming of Mbd3 and Gatad2a depleted, but not WT cells. We further characterized the codon combination that shows the highest variability, for each subset of ESPGs, CAPGs, or total differential genes (Fig. 6k,l). Surprisingly, the codon usage in ESPGs (red) and CAPGs (green) clustered at the lower and upper margins of the first principle component, respectively. The latter shows clear divergence in codon usage programs between the CAPGs and ESPGs: while ESPGs mainly tend to use codons that end with an G/C at the third codon position, CAP genes split into two programs: the genes that are induced during reprogramming, are encoded with A/T ending codons, while those that are repressed in the process mainly use G/C-ending codons (Fig. 6k,l, Extended Data Fig. 17a,b). Interestingly, we do not see any significant change in codon usage when comparing different time points of ESPGs, but we do see a very rapid and significant change in codon usage of CAPGs already emerging already between day0 and day1 (Fig. 6k) which underlies the global change observed during reprogramming (Fig. 6i). The efficiency of translation elongation is determined by the relation between the supply of tRNAs and the demand for specific tRNA types, governed by the representation of the 61 sense codons in the transcriptome. We thus asked whether the changes in codon usage along reprogramming are accompanied with a coordinated change in the tRNA pool. While methods for accurate estimation of tRNA expression are still developing^57,58^, it was shown that chromatin marks can serve as a proxy for tRNA expression^59^. We thus measured the chromatin mark H3K4me3 in the vicinity of the tRNA genes, and observed a change in the tRNA pool throughout reprogramming (Extended Data Fig. 16c). We next asked whether the change in the tRNA pool along reprogramming correspond to the observed change in the codon usage of the translated transcriptome. For this purpose, we calculated the expected translational efficiency for genes belonging to the most highly enriched GO categories corresponding to up/down regulated ESPGs and CAPGs (**see Methods**) based on their codon sequence and the tRNA epigenetic status. Remarkably, we observed a global significant positive correlation between the changes in transcription and translation, suggesting that the anticodons whose expression is elevated along reprogramming correspond to the codons that are enriched in the transcriptome of the respective cell state (Spearman r = 0.45, p< 4.5e-49, Extended Data Fig. 17d). However, while GO annotations that are associated with upregulated CAPGs showed an increase in translation efficiency (Fig. 6l), GO annotations associated with upregulated ESPGs show an opposite trend: a decrease in translation efficiency, corresponding to their G/C-ending codon preference (Fig. 6k-l, Extended Data Fig. 17a,b). The CAPG program is responsible for biosynthetic processes and is optimally regulated mainly by Myc and tRNA codon usage (Extended Data Fig. 18).

In summary, our experiments help better decipher the black box of reprogramming and provide a relatively more “continuous regulatory movie” of successful epigenetic reprogramming trajectory of mouse somatic cells to naïve pluripotency (Extended Data Fig. 18). In the future, increasing the dissection resolution by either integrating other layers of chromatin changes (e.g. Hi-C), higher frequency sampling during the 8 days reprogramming course or assaying deterministic reprogramming at the single cell level, will further facilitate unwinding the multilayered complexity of how cell states can be successfully and completely reconfigured.

## Author Contributions

A.Z., N.M, Y.R., N. N. and J.H.H conceived the idea for this project, designed and conducted experiments, and wrote the manuscript with contributions from other authors. L.W. generated and validated Tet1/2/3^flox/flox^ Mice. M.Z., R.M. and Y.R. conducted micro-injection and embryo-dissections. S.B. and S.G. conducted and supervised high-throughput sequencing. A.M. and Y.S.M. assisted in RNA-seq analysis. I.U. and H.H. conducted lncRNA analysis. W.J.G. and J.B. assisted in establishing and analyzing ATAC-seq experiments. E.C. assisted in DNA methylation sample preparation and analysis. H.G. and A.Z. performed tRNA codon usage analysis and experiment under the supervision of Y.P. I.A., D.A. and D.J. advised and assisted Hanna lab members in conducting ChIP-seq experiments and analysis. N.N. supervised all bioinformatics analysis and analyzed ChIP-seq data. S.P., M.A., I.M., S.H., A.A, J.B., D.S. and V.K. assisted in tissue culture and chromatin IP experiments. Y.S. generated and advised on conducting experiments involving RGM reporters. Y.R. and N.M engineered cell lines under S.V. supervision and designs. N.N. and J.H.H. supervised executions of experiments, adequate analysis of data and presentation of conclusions made in this project.

## Acknowledgements

J.H.H is supported by a generous gift from Ilana and Pascal Mantoux, and research grants from the: European Research Council, Flight Attendant Medical Research Council (FAMRI), Israel Science Foundation (ISF-ICORE, ISF-NFSC, ISF-INCPM & ISF-Morasha programs), Kamin-Yeda program, Minerva fund, Israel Cancer Research Fund (ICRF), Human Frontiers Science Program (HFSP), the Benoziyo Endowment fund, New York Stem Cell Foundation (NYSCF), Kimmel Innovator Research Award, the Helen and Martin Kimmel Institute for Stem Cell Research. J.H.H. is a New York Stem Cell Foundation (NYSCF)–Robertson Investigator. N.N. is supported by ISF-Morasha program. We acknowledge Y. Tabach for assistance on evolutionary conservation analysis; R. Jaenisch for RGM reporter lines and plasmids; Andreas Trump for Myc conditional mutant mESC lines. We thank Naama Barkai, Amos Tanay, Noam Stern-Ginossar, Tzachi Hagai and Bryce Carey for input and discussions during the course of conducting this project. We thank the Weizmann Institute management and board for providing critical financial and infrastructural support. In memory of Prof. Haim Garty who generously helped establish and fund the Hanna lab and this project.

## Methods

### Generation of Gatad2a-knockout Reprogrammable secondary MEF lines

Secondary MEF for Gatad2a^-/-^ cell line and WT-2 were obtained as described by Mor et al^15^. Shortly, iPSCs were established following primary reprogramming of cells using M2rtTA and TetO-OKSM-STEMCCA. The iPSC, harboring mCherry constitutive expression (to label viable cells) and ΔPE-GOF18-Oct4-GFP cassette (Addgene plasmid# 52382), were then subjected to CRISPR/Cas9 targeting Gatad2a (sgRNA-cgcctgatgtgattgtgct), resulting in Gatad2a-knockout cells (Extended Data Fig 1). Both Gatad2a-KO and its isogenic wild-type line (WT-2) were then injected into blastocysts, and MEF were harvested at E13.5. MEFs were harvested at E13.5 and grown in MEF medium, which contained 500 ml DMEM (Invitrogen), 10% fetal calf serum (Biological Industries), 1 mM glutamine (Invitrogen), 1% non-essential amino acids (Invitrogen), 1% penicillin– streptomycin (Invitrogen), 1% sodium pyruvate (Invitrogen). All animal studies were conducted according to the guideline and following approval by the Weizmann Institute IACUC (approval # 33550117-2 and 33520117-3). Cell sorting and FACS analysis were conducted on 4 lasers equipped FACS Aria III cells sorter (BD). Analysis was conducted with either DIVA software or Flowjo. Throughout this study, all cell lines were monthly checked for Mycoplasma contaminations (LONZA– MYCOALERT KIT), and all samples analyzed in this study were never tested positive or contaminated.

### Generation of reprogrammable Mbd3^flox/-^ secondary MEF lines

All secondary reprogrammable lines harbor constitutive expression of the M2rtTA from the Rosa26 locus and TetO OKSM cassette introduced either by viral transduction of knock-in in the Col1a1 locus. Secondary mouse embryonic fibroblast (MEF) from Mbd3^flox/-^ cell line (A12 clone: *Mbd3*^*flox/-*^ cell lines that carries the GOF18-Oct4-GFP transgenic reporter (complete *Oct4* enhancer region with distal and proximal enhancer elements) (Addgene plasmid #60527)) and WT-1 cell line (WT-1 clone that carries the deltaPE-GOF18-Oct4-GFP reporter (Addgene plasmid#52382) were previously described^13^. Note that we do not use Oct4–GFP or any other selection for cells before harvesting samples for conducting genomic experiments.

### Mouse embryo micromanipulation

Pluripotent mouse ESCs and iPSCs were injected into BDF2 diploid blastocysts, harvested from hormone primed BDF1 6 week old females. Microinjection into E3.5 blastocysts placed in M16 medium under mineral oil was done by a flat-tip microinjection pipette. A controlled number of 10-12 cells were injected into the blastocyst cavity. After injection, blastocysts were returned to KSOM media (Invitrogen) and placed at 37^o^C until transferred to recipient females. Ten to fifteen injected blastocysts were transferred to each uterine horn of 2.5 days post coitum pseudo-pregnant females.

### Reprogramming of MEF to naive ground state naive iPSC

Reprogramming of the optimally NuRD depleted and WT platform cell lines to IPSC was performed for the first 3 days with MES medium, which contained 500 ml DMEM (Invitrogen), 15% fetal calf serum, 1 mM glutamine (Invitrogen), 1% non-essential amino acids (Invitrogen), 1% penicillin– streptomycin (Invitrogen), 1% sodium pyruvate (Invitrogen), 0.1 mM *β*-mercaptoethanol (Sigma), 20 ng/ml human LIF (in house prepared). MES medium for reprogramming was supplemented with Doxycycline (DOX) (2 *μ*g ml-1), which activated the OKSM cassette and the reprogramming process. On day 3.5, medium was replaced to FBS-free media composed of: 500 ml DMEM (Invitrogen), 15% knockout serum replacement (Invitrogen; 10828), 1 mM glutamine (Invitrogen), 1% non-essential amino acids (Invitrogen), 0.1 mM *β*-mercaptoethanol (Sigma), 1% penicillin– streptomycin (Invitrogen), 1% sodium pyruvate (Invitrogen), 20 ng/ml recombinant human LIF (Peprotech or in house-prepared), CHIR99021 (3 *μ*M; Axon Medchem), PD0325901 (PD, 0.3-1*μ* M; Axon Medchem). After DOX treatment medium was replaced to KSR-based with the addition of MEK and GSK3 inhibitors (2i), supplemented with Doxycycline (DOX) (2 *μ*g ml-1). Cells were harvested at first time point (MEF) and every 24 hours until day 8, and were used for library preparation followed by sequencing. Cell was also collected for established iPSC line (after 3 passages or more), and control Mbd3^f/-^ or WT mouse ESCs. For all mouse iPSC reprogramming experiments, irradiated human foreskin fibroblasts were used as feeder cells, as any sequencing input originating from the use of human feeder cells cannot be aligned to the mouse genome and is therefore omitted from the analysis. All cell undergoing reprogramming were harvested without any prior passaging or sorting for any subpopulations during the reprogramming process. No blinding was conducted when testing outcome of reprogramming experiments.

### Primary and secondary reprogrammable lines by viral infection

For primary cell reprogramming, ~3x106 293T cells in a 10cm culture dish were transfected with jetPEI^®^ (Polyplus) 20ul reagent for 10ug DNA as follow: pPAX (3.5 *μ*g), pMDG (1.5 *μ*g) and 5*μ* g of the lentiviral target plasmid (pLM-mCerulean-cMyc (Plasmid #23244), FUW-STEMCCA-OKS-mCherry or FUW-M2rtTA, FUW-TetO-STEMCCA-OKS-mCherry (a gift kind from Gustavo Mostoslavsky). Viral supernatant was harvest 48 and 72 hours post transfection, filtered through 0.45micron sterile filters (Nalgene) and added freshly to the primary MEF that was isolated from Mbd3^flox/-^ chimeric mice (unless indicated otherwise). At day 4 cells was sorted by the relevant florescent filter (mCerulean (cMyc OE), mCherry (OSK OE) or double positive (OSK+M OE) cell was collected for RNA extraction or seeded for farther growth.

### Knockdown endogenous Myc during reprogramming

For secondary Mbd3^f/-^ OSK^2nd^ production, primary MEFs from Mbd3^flox/-^ chimeric mice were infected with FUW-TetO-STEMCCA-OKS-mCherry and FUW-M2rtTA. IPS cells were isolated and injected into BDF2 blastocysts for the isolation of secondary MEFs. Secondary MEFs were transfected at day −3 and again at day 0 (starting reprogramming by adding DOX) with siRNA for cMyc, lMyc, nMyc or control (Stealth siRNA-mix of 3 as indicated in the table below) with RNAiMAX (Invitrogen). For molecular analysis, cells were collected at day 3 and day 7 or day 8 as indicated.

|  |  |  |
| --- | --- | --- |
| Mouse-cMYC | Invitrogen | MSS-237326, MSS-237327, MSS-237328 |
| Mouse-lMYC | Invitrogen | MSS-275360, MSS-275361, MSS-275362 |
| Mouse-nMYC | Invitrogen | MSS-207081, MSS-207082, MSS-276042 |
| Stealth RNAi™<br>siRNA Negative Control,<br>Med GC | Invitrogen | 12935300 |

### Generation of triple Tet1,2,3^flox/flox^ mice and cell lines

Tet2^flox/flox^ mice were obtained from Jackson Laboratories (Stock number 017573). Tet1^flox/flox^ mice were generated by using conditional knockout targeting vector against Exon 4 in V6.5 ESC. After removal of Neomycin selection cassettes by Flippase in correctly targeted ESCs (validated both by Southern Blot and PCR analysis), chimeric blastocyst injections followed by successful germline transmission allowed us to establish Tet1^flox/flox^ mouse colony. Tet3^flox/flox^ mice were generated by gene targeting of the endogenous Exon 7 (contains Fe(ii) catalytic domain) Tet3 locus. After removal of Neomycin selection cassettes by Flippase in correctly targeted ESCs (validated both by Southern Blot and PCR analysis), chimeric blastocyst injections followed by successful germline transmission allowed us to establish Tet3^flox/flox^ mouse colony. Triple floxed homozygous mice were generated by interbreeding, after which Tet1^flox/flox^ Tet2^flox/flox^ Tet3^flox/flox^ mouse strain was obtained. Genotyping primers and strategy:

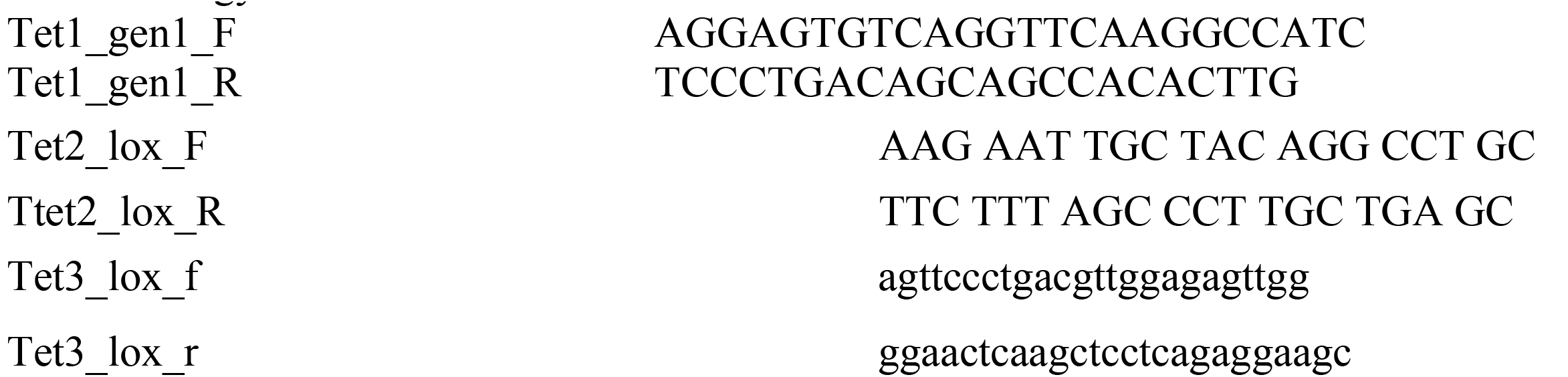

[The Tet1 floxed allele gives a band of 500bp, compared to the 450bp WT]

[The Tet2 flox allele gives a band of 427bp, compared to the 249bp WT].

[The Tet3 floxed allele gives a band of 300bp, compared to the 200bp WT].

MEFs, ESCs and iPSCs were derived from triple Tet1/2/3^flox/flox^ mice and were used as indicated in the figures. Deleting Gatad2a in Tet1/2/3^flox/flox^ iPSCs was done with CRISPR/Cas9 as indicated in Extended Data Fig. 1 and methods above.

### RT-PCR analysis

Total RNA was isolated using Trizol (ThermoFisher). 1 μg of DNase-I-treated RNA was reverse transcribed using a First Strand Synthesis kit (Invitrogen) and ultimately re-suspended in 100 *¼*l of water. Quantitative PCR analysis was performed in triplicate using 1/50 of the reverse transcription reaction in an Viia7 platform (Applied Biosystems). Error bars indicate standard deviation of triplicate measurements for each measurement.

### AP Staining

Alkaline phosphatase (AP) staining was performed with AP kit (Millipore SCR004) according to manufacturer’s instructions.

### Imaging, quantifications, and statistical analysis

Imaged were acquired with D1 inverted microscope (Carl Zeiss, Germany) equipped with DP73 camera (Olympus, Japan) or with Zeiss LSM 700 inverted confocal microscope (Carl Zeiss, Germany) equipped with 405nm, 488nm, 555nm and 635 solid state lasers, using a 20x Plan-Apochromat objective (NA 0.8). All images were acquired in sequential mode. For comparative analysis, all parameters during image acquisition were kept constant throughout each experiment. Images were processed with Zen blue 2011 software (Carl Zeiss, Germany), and Adobe Photoshop.

### ChIP-seq library preparation

Cells were crosslinked in formaldehyde (1% final concentration, 10 min at room temperature), and then quenched with glycine (5 min at room temperature). Fixed cells were lysed in 50 mM HEPES KOH pH 7.5, 140 mM NaCl, 1 mM EDTA, 10% glycerol, 0.5% NP-40 alternative, 0.25% Triton supplemented with protease inhibitor at 4 °C (Roche, 04693159001), centrifuged at 950*g* for 10 min and re-suspended in 0.2% SDS, 10 mM EDTA, 140 mM NaCl and 10 mM Tris-HCl. Cells were then fragmented with a Branson Sonifier (model S-450D) at −4 °C to size ranges between 200 and 800 bp, and precipitated by centrifugation. Antibody was pre-bound by incubating with Protein-G Dynabeads (Invitrogen 100-07D) in blocking buffer (PBS supplemented with 0.5% TWEEN and 0.5% BSA) for 1 h at room temperature. Washed beads were added to the chromatin lysate for an incubation period as detailed in the following table:

|  |  |  |
| --- | --- | --- |
| H3K4Me1 | Abcam ab8895 | 6h |
| H3K4Me3 | Abcam 8580 | 6h |
| H3K27ac | Abcam ab4729 | 6h |
| H3K27Me3 | Millipore 07-449 | 6h |
| H3K9Me3 | Abcam ab8898 | 6h |
| H3K36Me3 | Abcam ab9050 | 6h |
| H3K9Me2 | MBL MABI0317 | 6h |
| Oct4 | Santa Cruz SC8628 | Overnight (16h) |
| Klf4 | R&D AF3158 | Overnight (16h) |
| Sox2 | Millipore AB5603 | Overnight (16h) |
| C-Myc | Santa Cruz sc764 | Overnight (16h) |
| PolII | Santa Cruz sc899(N20) | Overnight (16h) |

Samples were washed five times with RIPA buffer, twice with RIPA buffer supplemented with 500 mM NaCl, twice with LiCl buffer (10 mM TE, 250mM LiCl, 0.5% NP-40, 0.5% DOC), once with TE (10Mm Tris-HCl pH 8.0, 1mM EDTA), and then eluted in 0.5% SDS, 300 mM NaCl, 5 mM EDTA, 10 mM Tris HCl pH 8.0. Eluate was incubated treated sequentially with RNaseA (Roche, 11119915001) for 30 min and proteinase K (NEB, P8102S) for 2 h in 65 °C for 8 h, and then. DNA was purified with The Agencourt AMPure XP system (Beckman Coulter Genomics, A63881). Libraries of cross-reversed ChIP DNA samples were prepared according to a modified version of the Illumina Genomic DNA protocol. All chromatin immunoprecipitation data are available at the National Center for Biotechnology Information Gene Expression Omnibus database under the series accession GEO no. GSE102518. [The following secure token has been created to allow review of record <u>**GSE102518**</u> while it remains in private status:]. Samples were run with various protocols and machines, for details see **Supplementary Table 1**.

### ChIP-seq analysis

**Alignment and peak detection:** We used bowtie2 software^60^ to align reads to mouse mm10 reference genome (UCSC, December 2011), with default parameters. We identified enriched intervals of all measured proteins using MACS version 1.4.2-1^61^. We used sequencing of whole-cell extract as control to define a background model. Duplicate reads aligned to the exact same location are excluded by MACS default configuration.

**TSS, TES and Enhancer definition:** Transcription start sites (TSS) and transcription end sites (TES) were taken from mm10 assembly (UCSC, December 2011). Promoters/TES intervals were defined as 1000bp around each TSS/TES, and enhancers were defined as 300bp around enhancer detection summit point (see enhancer identification below).

**Chromatin modification profile estimation in TSS, TES and in enhancers:** Chromatin modification coverage in the genomic intervals was calculated using in-house script. Shortly, the genomic interval is divided to 50bp size bins, and the coverage in each bin is estimated. Each bin is then converted to z-score by normalizing by the mean and standard deviation of the sample noise (Xˆj=(Xj-*¼*_noise)_/σ_noise_). Noise parameters were estimated for each sample from 6*10^7^ random bp across the genome. Finally, the 3^rd^ highest bin z-score of each interval is set to represent the coverage of that interval.

**Transcription factor binding in promoter and enhancer:** Promoter or enhancer was defined as bound by a TF if it overlapped a binding peak of the TF, as detected by MACS.

### PolyA-RNA-seq library preparation

Total RNA was isolated from indicated cell lines, RNA was extracted from Trizol pellets by Direct-zol RNA MiniPrep kit (Zymo), and utilized for RNA-Seq by TruSeq RNA Sample Preparation Kit v2 (Illumina) according to manufacturer’s instruction. See **Supplementary Table 1** for details of protocol and sequencing machine used.

### Small RNA-seq library preparation

1ug of total RNA from each sample was processed using the Truseq small RNA sample preparation kit (RS-200-0012 Illumina) followed by 12 cycles of PCR amplification. Libraries were evaluated by Qubit and TapeStation. For purification of the small RNA fragments, they were size selected using Blupippne machine (Sage Science) with 3% gel cassette followed by clean-up with minielute PCR purification kit (Qiagen). The libraries were constructed with different barcodes to allow multiplexing of 11 samples. See **Supplementary Table 1** for details of protocol and sequencing machine used.

### RNA-seq analysis: alignment, gene expression estimation

**Read Alignment for PolyA-RNA-seq** Tophat software version 2.0.10^62^ was used to align reads to mouse mm10 reference genome (UCSC, December 2011). FPKM values were calculated over all genes in mm10 assembly GTF (UCSC, December 2011), using cufflinks (version 2.2.1)^63^. Genes annotated as protein coding, pseudogene or lncRNA (n=24,439) were selected for further analysis.

**Read Alignment for Small RNA-seq** Bowtie software version 2 was used to align reads to mouse mm10 reference genome (UCSC, December 2011). FPKM values were calculated over all genes in mm10 assembly GTF (UCSC, December 2011), using cufflinks (version 2.2.1)^63^. Genes annotated as rRNA, miRNA, snoRNA were selected for further analysis.

Subsequently, PolyA and small RNA-seq FPKM were combined and processed together.

**Active and Differential genes** Gene was defined to be active in samples where FPKM is above 0.5 of the gene max value. Differential genes were defined by (FC>4) & (maximum value>1). Subsequent filtering was done to reject oscillatory or non-continuous time series by comparing he sum of derivatives to the total span. Specifically, the filtering scheme is 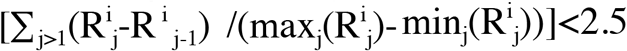, where j is the sample index, and i is the gene index.

**Expression HeatMap** Gene sorting in expression heat-maps (Fig. 1d) was done according to the average position of gene active samples, i.e. calculating the average of sample indexes (j) where the gene is active. **Unit normalized** FPKM was calculated using the following formula 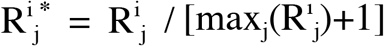 where j is the sample index, i is the gene index and FPKM=1 is the transcription noise threshold, and 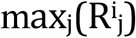 is the maximal level in each dataset. This normalization scheme allowed easy comparison of gene temporal patter with normalized dynamic range.

**Correlations** All correlation tests were done using Spearman correlation.

**PCA** PCA analysis (Fig. 1f) was carried out over all differential genes in unit normalization by Matlab (version R2011b) princomp command.

### Extended Differential lncRNA analysis

lncRNA dataset was annotated using PLAR^64^. FPKM values were calculated for all lncRNAs in PLAR mm9 dataset using cuffdiff (version 2.2.1)^63^. lncRNA coordinates were then converted to mm10 using liftOver utility. Differential lncRNAs were defined by (FC>4)&(maximum value>1). Subsequent filtering removed lncRNAs that were suspected to be expressed due to B1/B2-repeats by removing all sequence reads overlapping B1 or B2 repeats, resulting in 560 differential lncRNAs, out of them 221 differential lncRNA not previously annotated by Ensembl (Supplementary Table 2). Hierarchical clustering was performed over all differential lncRNAs with Spearman correlation metric and average linkage, further separating the differential lncRNAs to up, down regulated and intermediate induced lncRNAs (**Supplementary Table 3**).

RNA-seq data are available at the National Center for Biotechnology Information Gene Expression Omnibus database under the series accession GEO no. **<u>GSE102518</u>**. [The following secure token has been created to allow review of record GSE102518 while it remains in private status:].

### Phylogenetic analysis

Conservation scores of CAPGs and ESPGs were extracted from PhyloGene database^65^ http://genetics.mgh.harvard.edu/phylogene/.

### Functional Enrichment

Active genes at each sample (day) are tested for enrichment of functional gene sets taken from Gene Ontology (GO, http://www.geneontology.org), using Fisher exact test. Gene is defined to be active in samples where FPKM is above 0.5 of the gene max value. All enrichment values for each day were FDR corrected using Benjamini and Hochberg (1995) method^66^. GO annotations were filtered to include only annotations with FDR-corrected p-val<0.01 in at least two samples, annotations are sorted according to average position of enrichment pattern.

### Protein-DNA binding enrichment analysis

Active genes at each sample (day) are tested for enrichment (fisher exact test) to previously published protein-DNA binding ChIP-seq obtained from the Compendium, hmChip and BindDB databases. Gene is defined to be active in samples where RPKM is above 0.5 of the gene max value. All enrichment values for each day were passed through FDR test, using the Benjamini and Hochberg (1995) method. Subsequently, TF annotations per day were filtered to include only annotations with FDR-corrected Pval<10^-30^ in at least one sample. Further filtering the predicted TF to include only TF that are also differentially expressed during reprogramming according to our collected RNA-seq. The resulting predicted TF’s and their connectivity map from Compendium and hmChip are than merged where any connection exists in one of the databases also appears in the resulted connectivity matrix.

### ATAC-seq library preparation

Cells were trypsinized and counted, 50,000 cells were centrifuged at 500*g* for 3 min, followed by a wash using 50 *μ*l of cold PBS and centrifugation at 500*g* for 3 min. Cells were lysed using cold lysis buffer (10 mM Tris-HCl, pH 7.4, 10 mM NaCl, 3 mM MgCl2 and 0.1% IGEPAL CA-630). Immediately after lysis, nuclei were spun at 500*g* for 10 min using a refrigerated centrifuge. Next, the pellet was resuspended in the transposase reaction mix (25 *μ*l 2× TD buffer, 2.5 *μ*l transposase (Illumina) and 22.5 *μ*l nuclease-free water). The transposition reaction was carried out for 30 min at 37 °C and immediately put on ice. Directly afterwards, the sample was purified using a Qiagen MinElute kit. Following purification, the library fragments were amplified using custom Nextera PCR primers 1 and 2 for a total of 12 cycles. Following PCR amplification, the libraries were purified using a *Qiagen*MinElute Kit and sequenced as indicated in **Supplementary Table 1**.

### ATAC-seq analysis

Reads were aligned to mm10 mouse genome using Bowtie2 with the parameter -X2000 (allowing fragments up to 2 kb to align). Duplicated aligned reads were removed using Picard MarkDuplicates tool with the command REMOVE_DUPLICATES=true. To identify chromatin accessibility signal we considered only short reads (≤ 100bp) that correspond to nucleosome free region.

### Identifying accessible chromatin regions

To detect and separate accessible loci in each sample, we used MACS version 1.4.2-1 with–call-subpeaks flag (PeakSplitter version 1.0). Next, summits in previously annotated spurious regions were filtered out using a custom blacklist targeted at mitochondrial homologues. To develop this blacklist, we generated 10,000,000 synthetic 34mer reads derived from the mitochondrial genome. After mapping and peak calling of these synthetic reads we found 28 high-signal peaks for the mm10 genome. For all subsequent analysis, we discarded peaks falling within these regions.

### Enhancer Identification

Each ATAC-seq peak in each sample was represented by a 300bp region around the summit center. H3K27ac peaks were detected in a similar manner, using MACS version 1.4.2-1, and merged for all time points using bedtools merge command. All ATAC peaks were filtered to include only peaks which co-localized with the merged H3K27ac peaks, meaning only ATAC peaks that have H3K27ac mark on at least one of the time points were passed to further processing. Finally, the peaks from all samples were unified and merged (using bedtools unionbedg and merge commands), further filtered to reject peaks that co-localized with promoter or exon regions based on mm10 assembly (UCSC, December 2011). Finally we were left with 93,137 genomic intervals which we annotated as active enhancers^67^, of which 78% of overlap with H3K4me1 modification, and 69% are bound by at least one of the transcription factors mapped (RNA PolII/O/S/K/M) (Extended Data Fig. 3a). All enhancers were then annotated by their most proximal gene using annotatePeaks function (homer/4.7 package). Enhancers were considered as differential if both their ATAC-seq and H3K27ac signals show significant change during reprogramming (min zscore<0.5, max zscore>1.5, for both chromatin marks). ATAC-seq data are deposited under GEO no. **<u>GSE102518</u>**. [The following secure token has been created to allow review of record GSE102518 while it remains in private status:]

### Generating ATAC-seq normalized profiles in TSS and in enhancers

ATAC-seq profiles were calculated using in-house script over all genomic intervals defined for TSS and enhancers. Shortly, the genomic interval is divided to 50bp size bins, and the coverage in each bin is estimated. Each bin is then converted to z-score by normalizing each position by the mean and standard deviation of the sample noise (Xˆj=(Xj-µ_noise)_/σ_noise_). Noise parameters were estimated for each sample from 6*10^7^ random bp across the genome. Finally, the 3^rd^ highest bin z-score of each interval is set to represent the coverage of that interval.

### Whole-Genome Bisulfite Sequencing (WGBS) library preparation

DNA was isolated from snap-frozen cells using the Quick-gDNA mini prep kit (*Zymo*). DNA was then converted by bisulfite using the EZ DNA Methylation-Gold kit (*Zymo*). Sequencing libraries were created using the EpiGnome Methyl-Seq (*Epicentre*) and sequenced as indicated at **Supplementary Table 1**

### Reduced-Representation Bisulfite (RRBS) library preparation

RRBS libraries were generated as described previously with slight modifications^40^. Briefly, DNA was isolated from snap-frozen cell pellets using the Quick-gDNA mini prep kit (Zymo). Isolated DNA was then subjected to MspI digestion (NEB), followed by end repair using T4 PNK/T4 DNA polymerase mix (NEB), A-tailing using Klenow fragment (3′5′ exo-) (NEB), size selection for fragments shorter than 500 bp using SPRI beads (Beckman Coulter) and ligation into a plasmid using quick T4 DNA ligase (NEB). Plasmids were treated with sodium bisulphite using the EZ DNA Methylation-Gold kit (Zymo) and the product was PCR amplified using GoTaq Hot Start DNA polymerase (Promega). The PCR products were A-tailed using Klenow fragment, ligated to indexed Illumina adapters using quick T4 DNA ligase and PCR amplified using GoTaq DNA polymerase. The libraries were then size-selected to 200–500 bp by extended gel electrophoresis using NuSieve 3:1 agarose (Lonza) and gel extraction (Qiagen). See **Supp Table 1** for sequencing protocol used.

## Methylation Analysis of WGBS and RRBS

### Alignment of RRBS data

The sequencing reads were aligned to the mouse mm10 reference genome (UCSC, December 2011), using Bismark aligner^68^ (parameters -n 1 -l 20). Mapping was done independently for the two ends of each pair. Read pairs that mapped uniquely to two different fragments were discarded. In cases where one read uniquely mapped on a restriction site but its pair could not be mapped uniquely or could not be mapped at all, we attempted to re-align the entire read pair to the fragment. Read pairs showing more than one unconverted non-CpG cytosine, which occur at very low frequency were filtered out.

### Alignment of WGBS data

The sequencing reads were aligned to the mouse mm10 reference genome (UCSC, December 2011), using a proprietary script based on Bowtie2. In cases where the two reads were not aligned in a concordant manner, the reads were discarded.

### Methylation estimation

Methylation levels of CpGs calculated by RRBS and WGBS were unified. Mean methylation was calculated for each CpG that was covered by at least 5 distinct reads (X5). Average methylation level in various genomic intervals was calculating by taking the average over all covered X5 covered CpG sites in that interval.

### Correlation of chromatin modifications

Correlation between chromatin modification to gene expression and to accessibility signal were estimated using Spearman correlation (Figure 4a, 5a, Extended Data Fig 5b-c). Promoters or enhancers with z-score above zero were included in the analysis, resulting in different number of promoter or enhancers for each chromatin marks (which are indicated in the figures).

### Cross-correlation of chromatin modifications

Cross correlation method^50,51^ measures the overlap between two signals, while shifting the signals in their x-axis (convolution). In our case, the x-axis is time. Cross correlation score was calculated using Matlab R2013b xcorr command. The offset showing the highest xcorr coefficient was defined as the optimal offset between the two signals. Cross-correlation was calculated in three systems: (i) Between chromatin modifications in promoters and gene expression pattern of ESPGs (Figure 5c). (ii) Between chromatin modifications and accessibility signal in differential enhancers (Figure 5f). (iii) Between chromatin modifications in promoters and enhancers that are associated with these promoters, and gene expression (Extended Data Fig. 15a).

In all these cases promoters/enhancers were included only if the modification z-score was changing (max-min>0.5), resulting in different number of promoters/enhancers as indicated in the graphs.

### Combinatorial analysis for histone marks localization

To quantify all possible combinations of epigenetic modifications (Figure 5b, 5e, Extended Data Fig 13f), we transformed our epigenetic data to a binary code in each genomic region (promoter/enhancer). Each epigenetic mark in promoter or enhancer was considered high (value=1) if its z-score was above 1.5. For each sample, the percentage of each combination is presented. Combinations which are less than 3% of the total combinations in every sample are presented as “other” (gray color).

### Motif analysis

Enriched binding motifs were searched in various genomic intervals (Extended Data Fig. 6,7a) using findMotifsGenome function from homer software package version 4.7^69^, using the software default parameters.

### Motif analysis in open vs. closed binding targets

In order to find binding motifs in open vs. closed binding targets (Fig. 2e, Extended Data Fig. 7c) we followed the analysis outline presented by Soufi et al^31^: We considered binding peaks of O/S/K/M in Day1, identified by MACS as explained. We calculated nucleosome occupancy in a 200bp window in the summit of the peak, and in two 100bp flanking regions on the two sides of the central window. Nucleosome occupancy was estimated from ATAC-seq data, measured in Day1, using nucleoatac occ software^70^. Top 2000 binding sites with highest center/flanking ratio were selected as closed sites (as long as ratio >1), and bottom 2000 sites were selected as open sites (as long as ratio <1). Next, motif search and annotation was done as in Soufi et al^31^, using DREME, Centrimo and TOMTOM software, of MEME suit^71^.

**Box plot analysis** Box-plots show 25-th and 75-th percentile of the represented distribution values, with median marked by the mid-line. The whiskers extend to the most extreme data point which is no more than 1.5 times the interquartile range from the box, and outliers are not presented.

## Translation Analysis

### Coding sequences

The coding sequences of *M. musculus* were downloaded from the Consensus CDS (CCDS) project (ftp://ftp.ncbi.nlm.nih.gov/pub/CCDS/).

### tRNA gene copy numbers

The tRNA gene copy numbers of *M. musculus* were downloaded from the Genomic tRNA Database (http://lowelab.ucsc.edu/GtRNAdb/) (Lowe and Eddy, 1997).

### Estimating translational efficiency by chromatin modification signature in the vicinity of tRNAs

We estimated translation efficiency of genes using the ‘‘tRNA activation index’’ (tACI) which was introduced previously by Gingold *et al*.^58^. This measure is calculated similarly to the tRNA Adaptation Index (tAI) measure of translation efficiency^72^, with one change—tRNA availabilities are determined based on chromatin modification in the vicinity of the tRNA genes rather than by gene copy numbers. Specifically, we set the activation score of each individual tRNA gene to be the maximal read per megabase (RPM) value of the activation-associated modification H3K4me3 across a region spanning the 500 nucleotides upstream to the first nucleotide of the mature tRNA. Individual tRNA genes, for which no signal enrichment was found, were classified as ‘‘not activated.’’ Next, we defined the activation score of each tRNA type (anticodon) by the sum of the activation scores of its gene copies. Then, we determined the translation efficiency of each of the 61 codon types by the extent of activation of the tRNAs that serve in translating it, incorporating both the fully matched tRNA as well as tRNAs that contribute to translation through wobble rules^73^.

Formally, the translation efficiency score for the *i*–th codon is

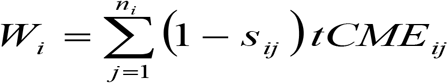

where *n* is the number of types of tRNA isoacceptors that recognize the *i*-th codon, *tCME*_*ij*_denotes the sum of the chromatin modification scores of the activated copies of the *j*-th tRNA that recognizes the *i*-th codon, and *S*_*ij*_ corresponds to the wobble interaction, or selective constraint on the efficiency of the pairing between codon *i* and anticodon *j*, as was determined and implemented for the original tAI measure. As done in the original tAI formalism by dos Reis et al., the scores of the 61 codons are further divided by the maximal score (yielding w_i_ as the normalized scores for each codon type), and finally, the tACI value of a gene with L codons is then calculated as the geometric mean of the *w*_*i*_’s of its codons

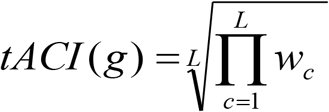

## Data Availability

All RNA-seq, ATAC-seq, ChIP-seq and methylation data are available to download from NCBI GEO, under super-series GSE102518.

## Extended Data Figure Legends

**Extended Data Fig. 1:**
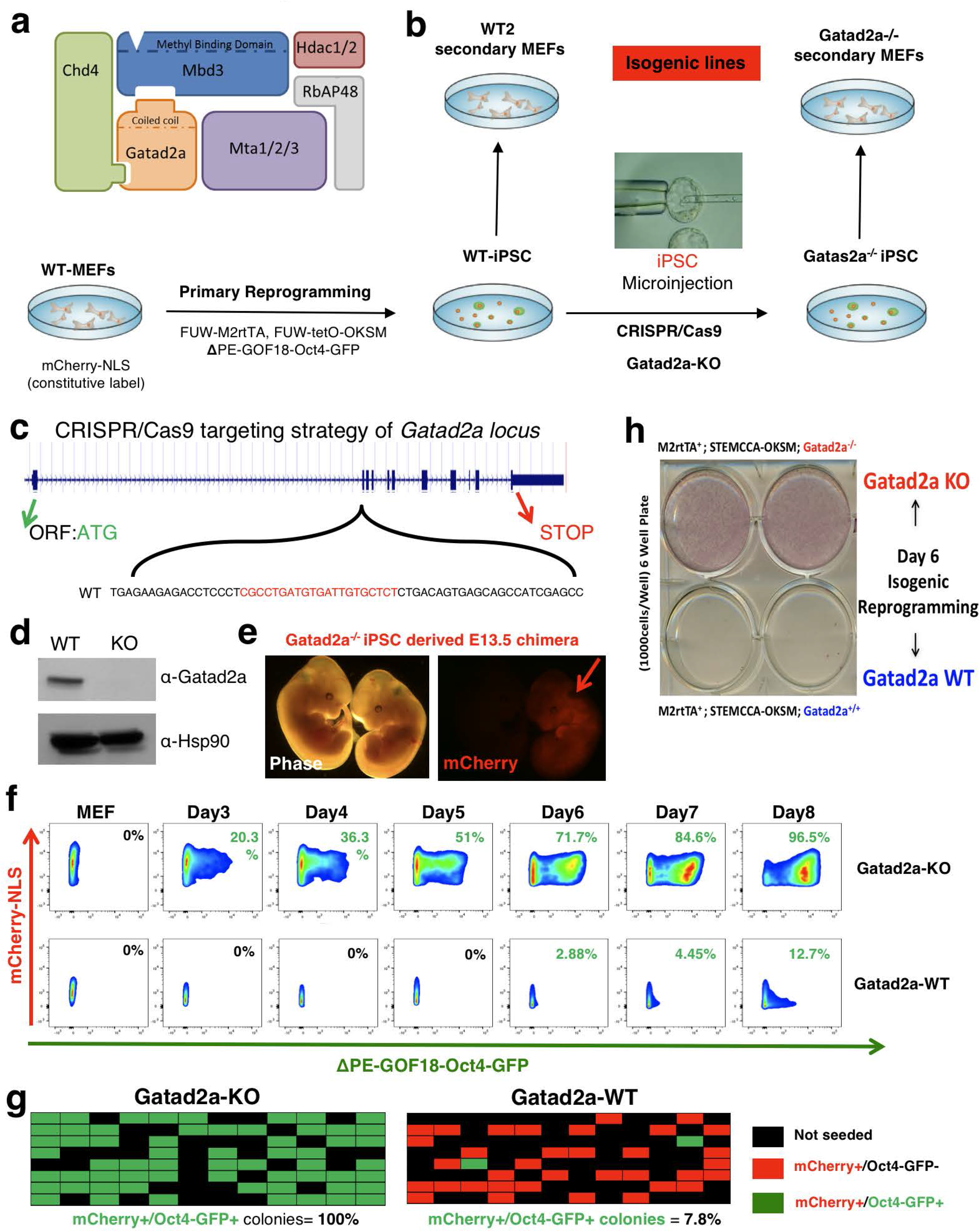
Neutralizing Gatad2a, a NuRD specific component, facilitates deterministic and highly efficient naïve iPSC induction from murine MEFs. a. Scheme depicting known components of Mbd3/NuRD complex. Gatad2a (also known as P66a) is a poorly characterized NuRD specific component. b. In order to generate WT and Gatad2a depleted isogenic reprogrammable systems and MEFs, the following strategy was applied: E13.5 WT MEFs constitutively labeled with nuclear-mCherry transgene, were reprogrammed to pluripotency using lentiviral transduction with TetO-OKSM-STEMMCA and M2rtTA constructs. Primary iPSC clone was established and transduced with naïve pluripotency specific ΔPE-GOF18-Oct4-GFP transgene. Sub cloned WT-2 line was validated for specificity of ΔPE-GOF18-Oct4-GFP reporter (homogenously turned on in 2i/LIF and FBS/LIF naïve conditions, and shut down upon differentiation). WT-2 iPSC line was then subjected to CRISPR/Cas9 targeting to generate Gatad2a knockout. A validated KO iPSC lines were then injected to blastocysts and returned to foster-mothers, and secondary reprogrammable MEFs were extracted from embryos at E13.5. c. CRISPR/Cas9 targeting strategy of murine Gatad2a locus. d. Western blot validation of Gatad2a knockout in iPSC harboring M2rtTA and TetO-OKSM cassettes e. Representative images of Gatad2a^-/-^ iPSC derived E13.5 chimera. Red arrow highlights mCherry+ chimera which originates from mCherry labeled iPSCs that were microinjected. f. Representative flow cytometry measurements of ΔPE-GOF18-Oct4-GFP reactivation dynamics in polyclonal/bulk Gatad2a-WT and Gatad2a-KO isogenic cell lines. Reprogramming was conducted as indicated in Fig. 1a. on human irradiated fibroblasts (used as feeder cells). Throughout the course of the reprogramming experiment the cells were not passaged to avoid any biases. g. Reprogramming of secondary MEFs seeded as single cells. A representative summary of single-cell experiment. Secondary isogenic Gtad2a WT and KO reprogrammable MEFs carrying constitutively expressed mCherry-NLS and naïve pluripotency specific ΔPE-GOF18-Oct4-GFP reporter were sorted and seeded as single-cell. Reprogramming was initiated by Dox administration according to Fig. 1a. Reprogramming efficiency was assessed after 8 days based on the number of wells in which mCherry+ cells formed an Oct4-GFP positive colony. Throughout the course of the reprogramming experiment the cells were not passaged to avoid any biases. h. Bulk iPSC reprogramming as in f., but experiment was terminated after 6 days and iPSC colony formation was evaluated by Alkaline Phosphatase staining (AP+). We have also found that complete inhibition of Gatad2a (also known as P66a), a NuRD specific subunit, does not compromise somatic cell proliferation as previously seen upon complete Mbd3 protein elimination, and yet disrupts Mbd3/NuRD repressive activity on the pluripotent circuitry and yields 90-100% highly-efficient reprogramming within 8 days as similarly observed in Mbd3 hypomorphic donor somatic cells (Extended Data Fig. 1 and Mor N. et al.^15^ *Under final preparation*).

**Extended Data Fig. 2:**
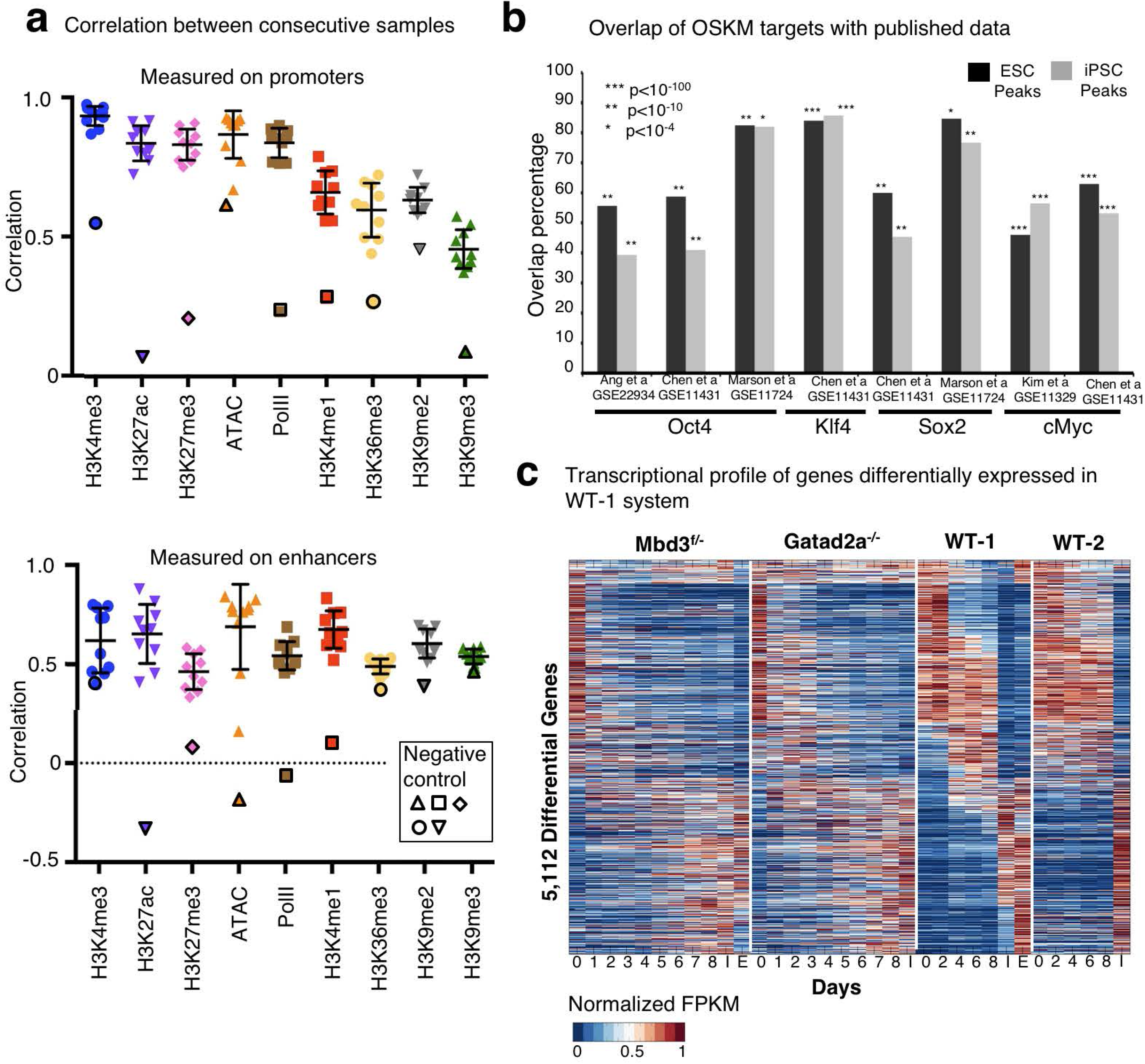
Epigenetic and transcriptional measurements are reproducible across highly efficient systems used. a. Correlation between consecutive samples in Mbd3^f/-^ system (MEF-day1, day1-day2, day2-day3… day8-IPS), measured over all ESPGs promoters (promoters with differential chromatin pattern, n=3,593, top), or all differential enhancers (n=40,174, bottom), for each chromatin mark. Negative controls were calculated between MEF and IPS, are marked with solid border. b. Overlap between binding targets of Oct4, Sox2, Klf4 or Myc, and previously published binding data of the same factors, calculated in ES and IPS samples. Percentage out of our measured binding targets is presented, along Fisher exact test p-values. c. Transcriptional profiles of Mbd3^f/-^ and Gatad2a^-/-^ reprogramming systems are nearly indistinguishable. Global transcriptional pattern of 5112 differential genes in WT-1 system, sorted by their temporal pattern in WT-1 system. HeatMap represents unit-transformation of FPKM values (see Methods).

**Extended Data Fig. 3:**
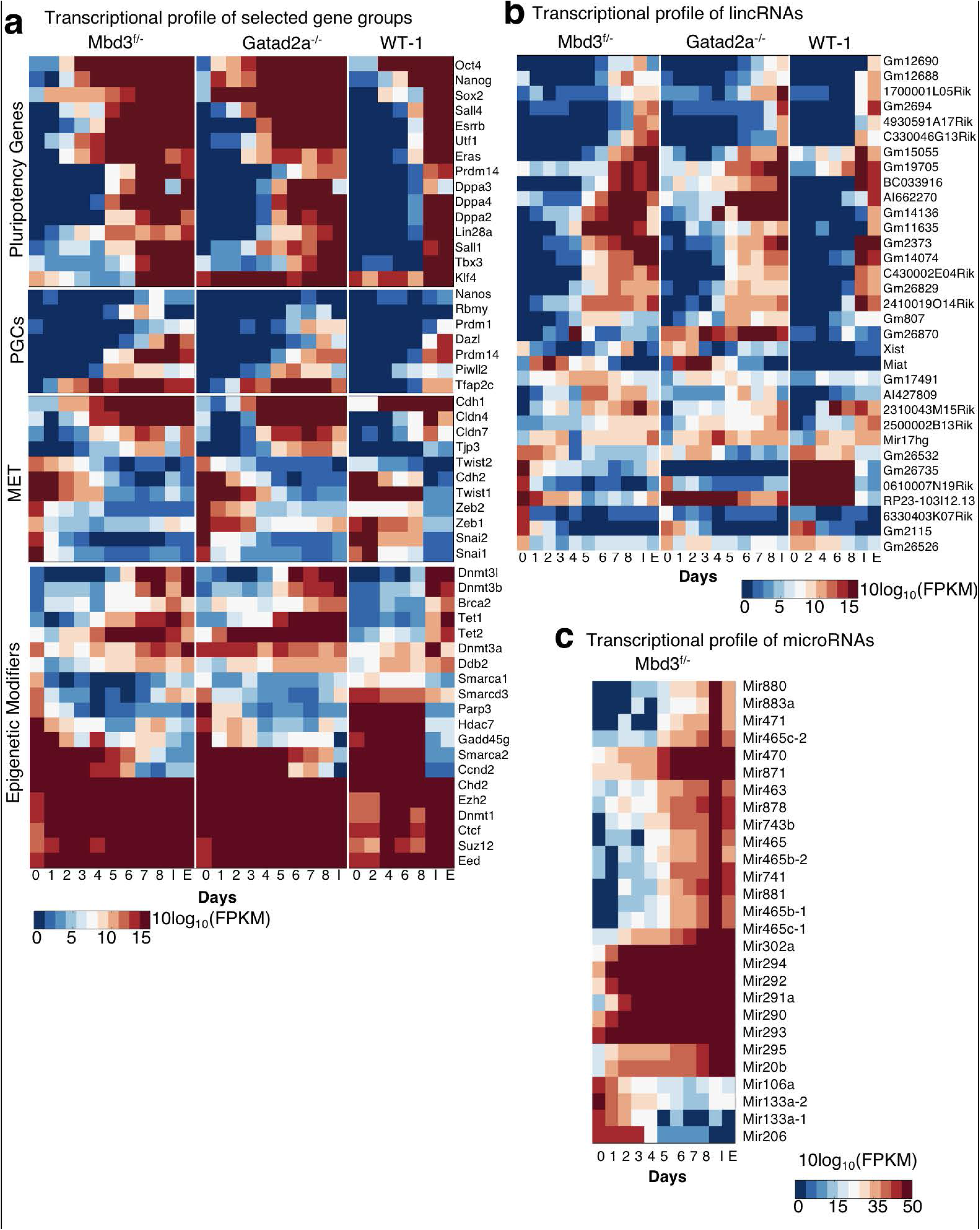
Continuous changes in specific gene groups of interest during the course of conducive naïve iPSC reprogramming. a. Transcription heatmap of Pluripotent genes, Primordial germ cell genes (PGCs), Mesenchymal to epithelial Transition (MET), and Epigenetic modifiers. Log_10_(FPKM) values are presented in Mbd3^f/-^, Gatad2a^-/-^ and WT-1 systems. b. Transcription heatmap of differential lncRNAs, annotated according to mm10 (UCSC, December 2011). Log_10_(FPKM) values are presented. c. Transcription heatmap of differential microRNAs (FC>600), measured using small-seq assay in Mbd3^f/-^ system.

**Extended Data Fig. 4:**
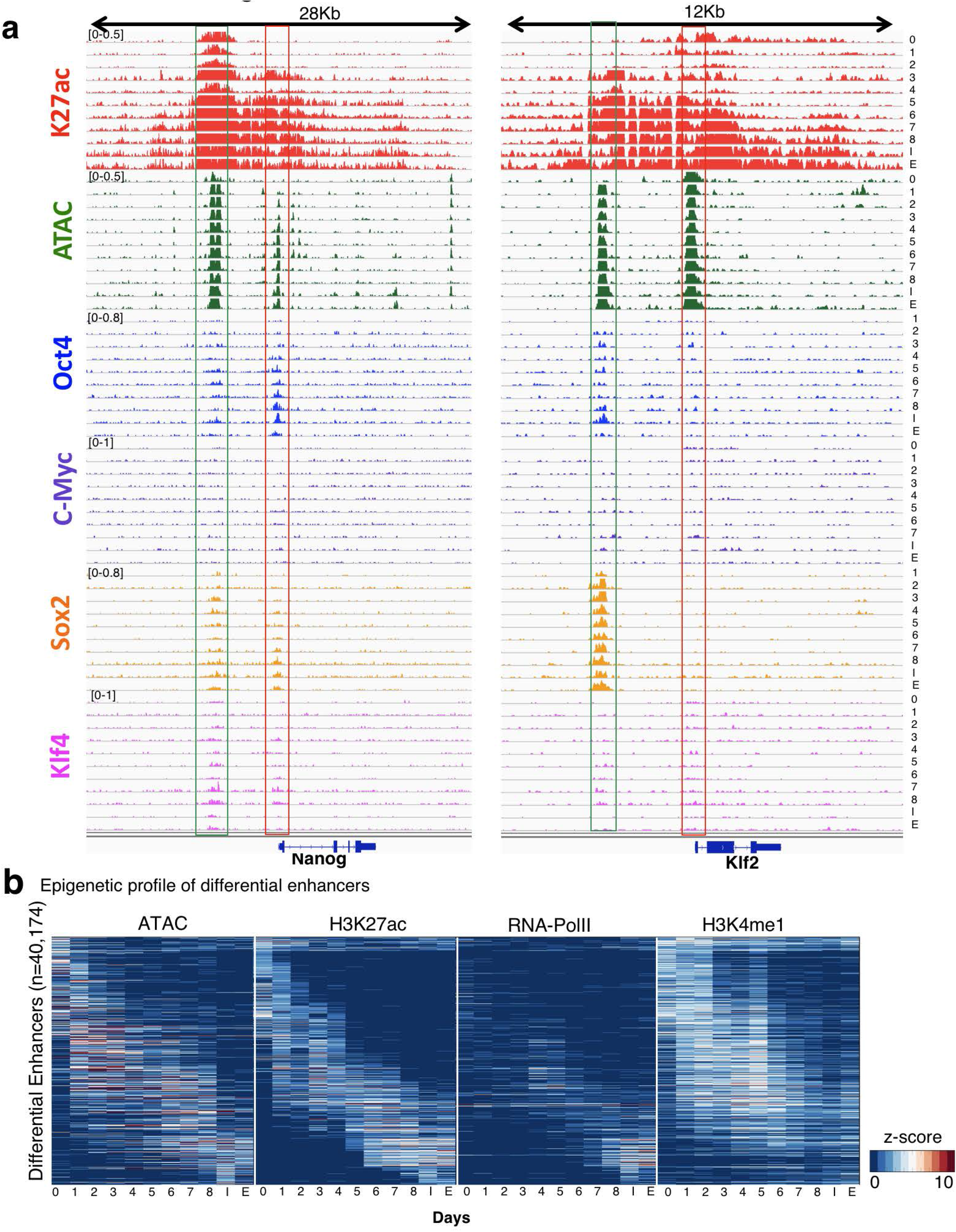
Massive epigenetic changes in enhancers detected during reprogramming. a. Landscape around two representative gene examples (Nanog and Klf2, promoters in red box), showing their associated enhancers (green box). Signals are normalized to sample size (RPM), and IGV data range is indicated on top left corner of the signal. b. Profiles of 40,174 differential enhancers along reprogramming, showing H3K4me1, H3K27ac, DNA accessibility and PolII binding.

**Extended Data Fig. 5:**
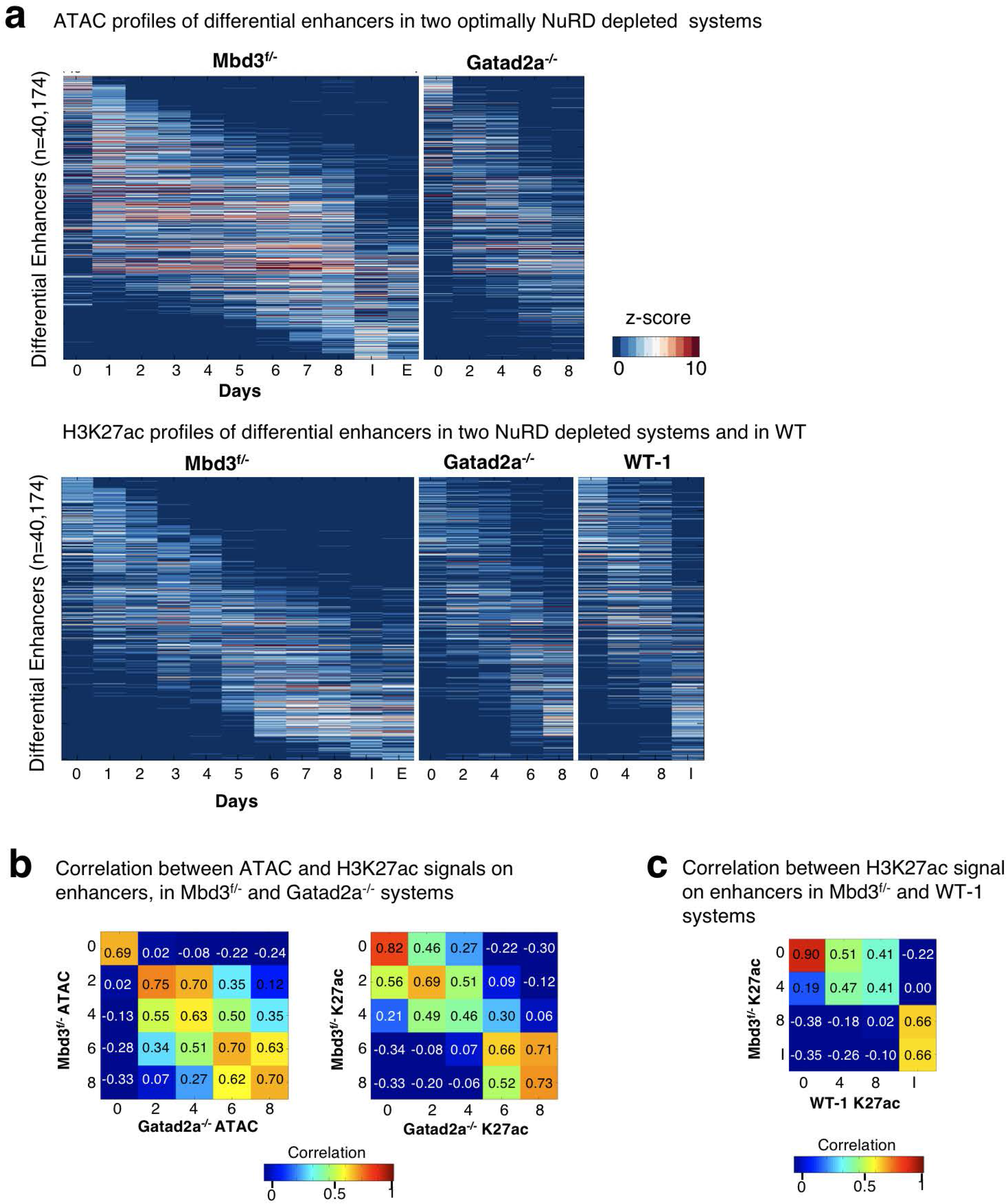
Coordinated and rapid epigenetic change in enhancers detected in two optimally NuRD-depleted systems. a. DNA accessibility (Top) and H3K27ac (bottoms) signals of 40,174 differential enhancers (rows), in Mbd3^f/-^, Gatad2a^-/-^, and WT-1 systems. b. Spearman correlation matrix comparing DNA accessibility (left) and H3K27ac (right) measured on enhancers in Mbd3^f/-^ and Gatad2a^-/-^ systems. c. Spearman correlation matrix comparing H3K27ac measured on enhancers in Mbd3^f/-^and WT systems. H3K27ac enhancer profile in WT-1 day8 is more similar to Mbd3^f/-^ day4, than to day8.

**Extended Data Fig. 6:**
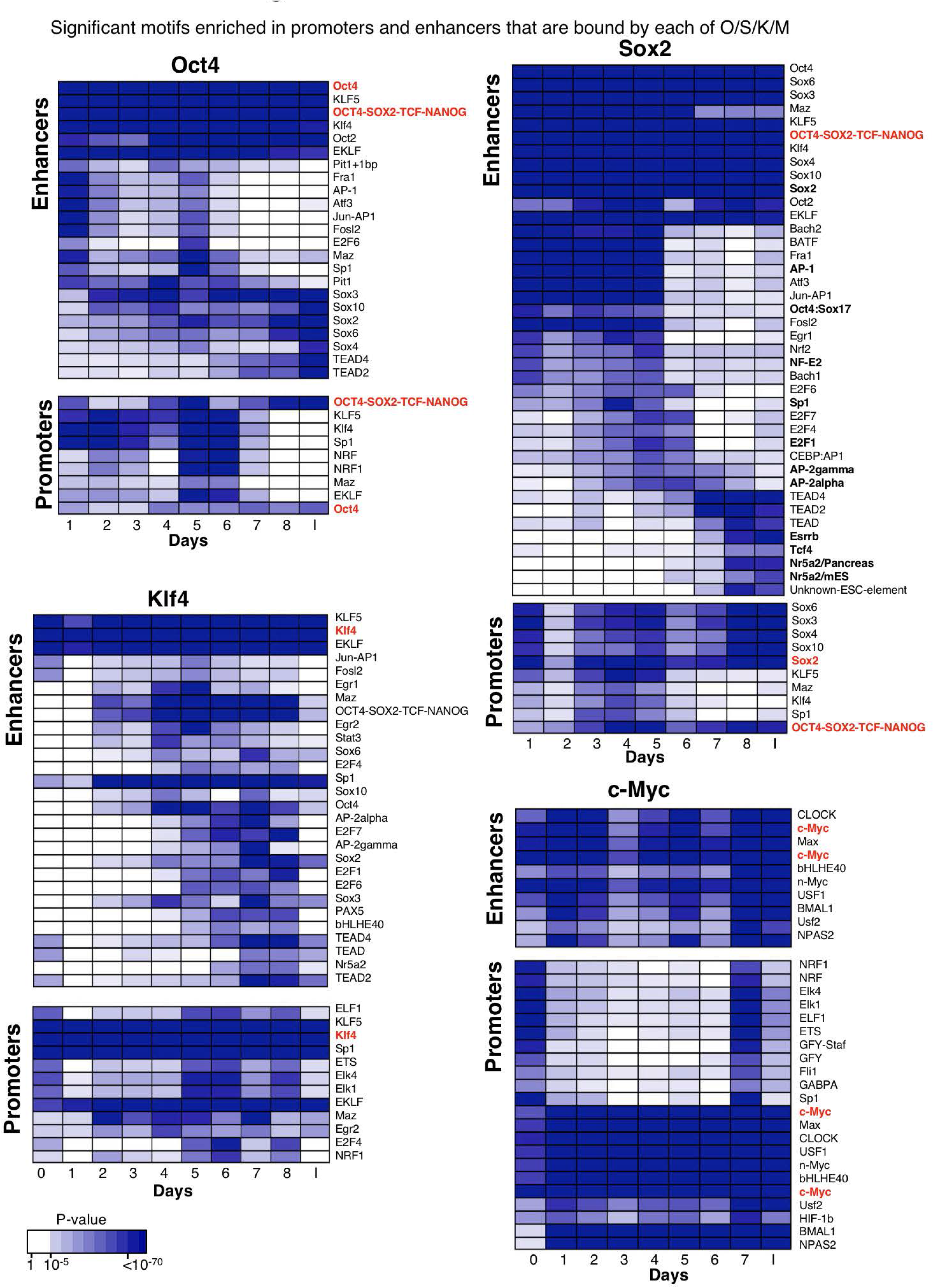
Dynamic changes in motif enrichment in OSKM-bound enhancers during conducive reprogramming trajectory. Significant motifs enriched in promoters and enhancers that are bound by each of Oct4, Sox2, Klf4 and c-Myc at different days of reprogramming, as detected by Homer/4.7 software^69^. P-values, indicated by color shade, were reported by Homer, and are FDR corrected. Motifs which are significantly enriched (corrected p<10^-30^) in at least one time point are presented.

**Extended Data Fig. 7:**
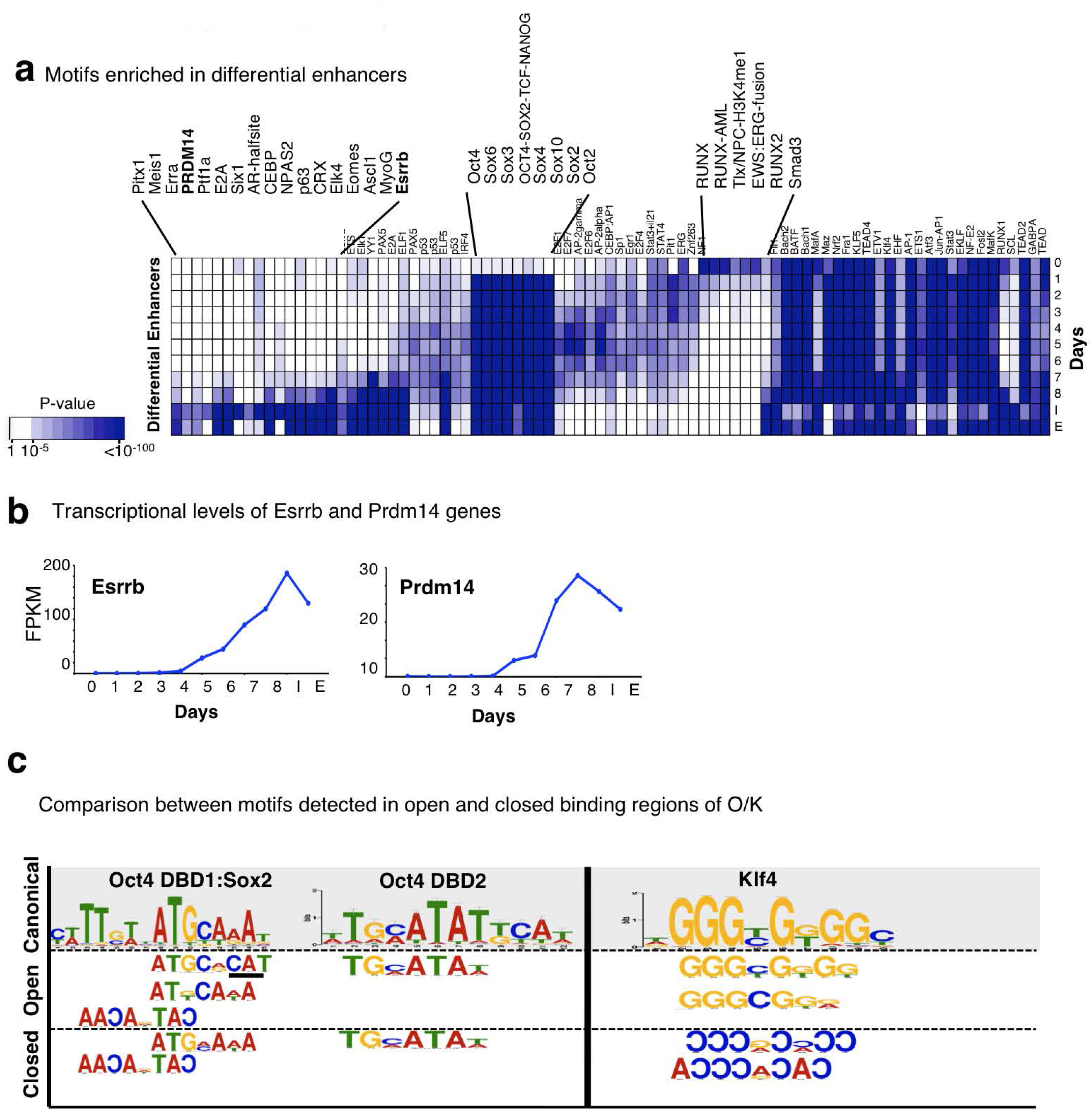
Stage-specific enhancer binding analysis show low affinity of OSKM to MEF-specific enhancers. a. Motifs enriched in differential enhancers that are active in each day of reprogramming (ATAC-seq z-score > 1.5). Motifs were detected by Homer/4.7 software^69^. P-values, indicated by color shade, were reported by Homer, and are FDR corrected. Motifs which are significantly enriched (corrected p<10^-50^) in at least one-time point are presented. b. Expression levels of Esrrb and Prdm14 transcription factors during conducive reprogramming. c. Motifs found in “closed” vs. “open” binding targets of the indicated transcription factor. Accessibility of targets was calculated based on ATAC-seq (See Methods). Motifs found in OSK binding targets calculated in Mbd3^f/-^ day1. Motifs that are different between open and closed binding targets are marked in black line. Complementary motifs to canonical motif appear in reverse order.

**Extended Data Fig. 8:**
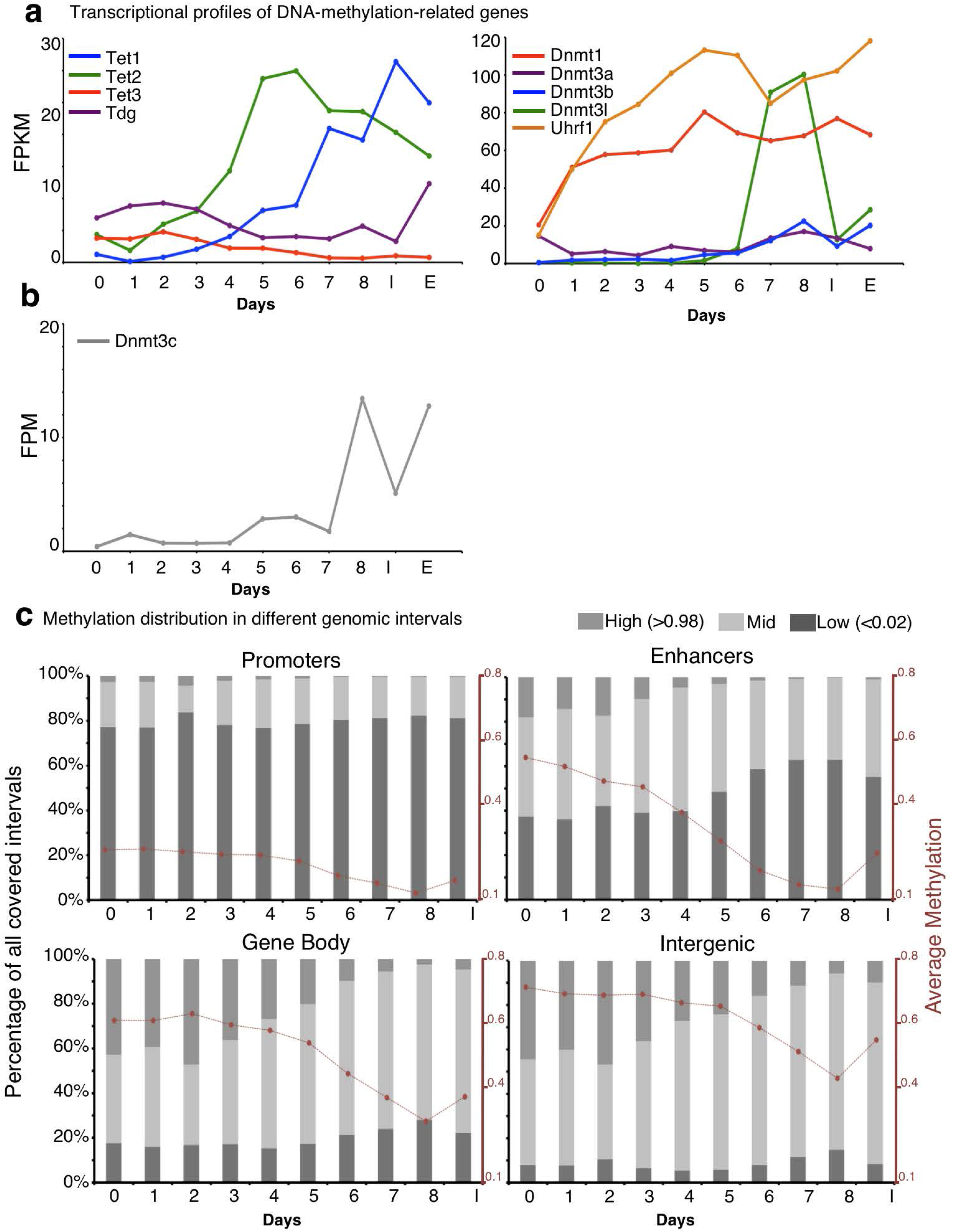
Global DNA demethylation and expression changes in Tet and Dnmt enzymes observed during reprogramming. a. Expression level of genes related to DNA methylation and demethylation. Expression levels are indicated as FPKM. b. Expression level of newly identified Dnmt3c gene^34^. Expression levels are indicated as FPM. c. Distribution of low (<0.02), mid (0.02-0.98) and high (>0.98) methylation in four genomic regions: Promoters (1000bp around the TSS), Enhancers, Gene body (TSS to TES) and Intergenic regions (1000bp tiles), calculated in each day of reprogramming.

**Extended Data Fig. 9:**
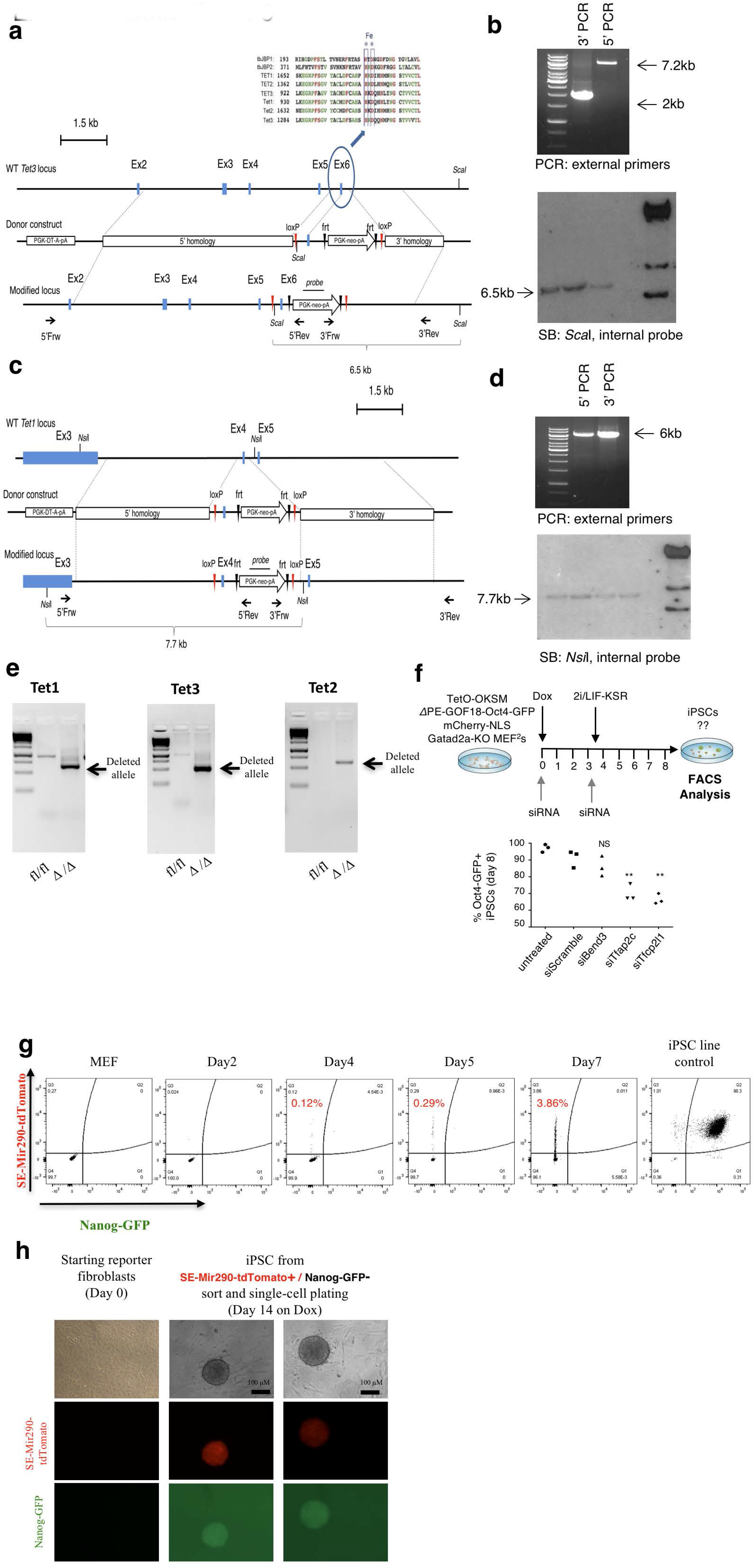
DNA demethylation is essential for rapid and deterministic reprogramming and is an early predictor for successful reprogramming. a. Targeting scheme for generating Tet3 conditional knock-out mouse model. b. PCR and Southern blot confirmation of correctly targeted clones. c. Targeting scheme for generating Tet1 conditional knock-out mouse model. d. PCR and Southern blot confirmation of correctly targeted clones. e. Genotyping validation example of Tet1/Tet2/Tet3 triple-conditional knock-out MEF cells, following Cre-Recombinase treatment. f. Reprogramming efficiency (measured by Oct4 GFP+ cells percentage) was measured after 8 reprogramming days of Gatad2a-KO secondary MEF, following siRNA treatments targeting Tfcp2l and Tfap2c, on days 0 and 3. Both treatments resulted in 20-30% reduction of reprogramming efficiency comparing to control specimen (siScramble). **p<0.01, ***p<0.001, Student t-test. n=3, error bars indicate SD. g. Flow cytometry measurements of both Nanog-GFP reactivation and SE-Mir290-tdTomato during reprogramming of secondary fibroblasts. Sorting for RGM-Mir290-tdTomato+/Nanog-GFP-cells and double negative cells was done at day 5. Please note that single positive Nanog-GFP+ cells do not exist in naive reprogramming conditions as applied herein. h. Representative pictures of iPSC colonies obtained at day 14, originating following single cell sorting of Nanog-GFP-/RGM-SE-Mir290-tdTomato cells at day 4 followed by continued DOX treatment. Donor fibroblast prior to DOX treatment (Day 0) are shown as negative controls. Scale bars = 100µM.

**Extended Data Fig. 10:**
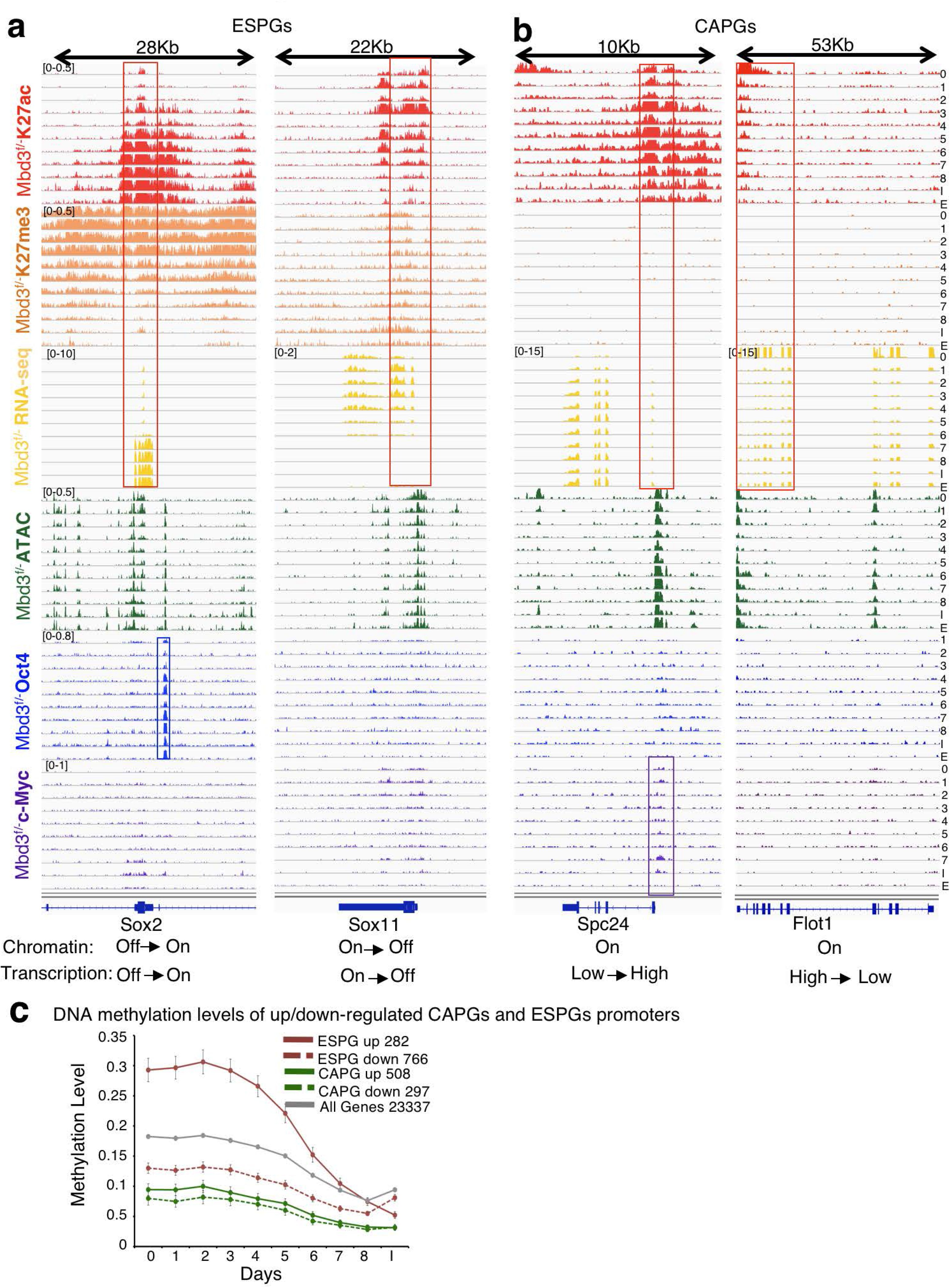
Different epigenetic landscape shown in ESP and CAP gene promoters and enhancers. a. ChIP-seq landscape showing binding of H3K27ac, H3K27me3, Oct4, c-Myc, alongside ATAC-seq and RNA-seq. Example of two genes with “epigenetically switched promoters” (ESPGs). Note that the change in H3K27ac and H3K27me3 corresponds to the change in expression (red box). Oct4 enhancer binding is highlighted in blue box b. Example of two genes with “consistently active promoters” (CAPGs). Note the constitutive high level of H3K27ac and low level of H3K27me3 regardless to the change in expression (red box). cMyc promoter binding is highlighted in purple box. Signals are normalized to sample size (RPM), and IGV data range is indicated on top left corner of the signal. c. DNA Methylation level of upregulated and downregulated ESPGs (red) and CAPGs (green), compared to all genes (gray), showing that CAPGs are hypomethylated, regardless of their expression pattern.

**Extended Data Fig. 11:**
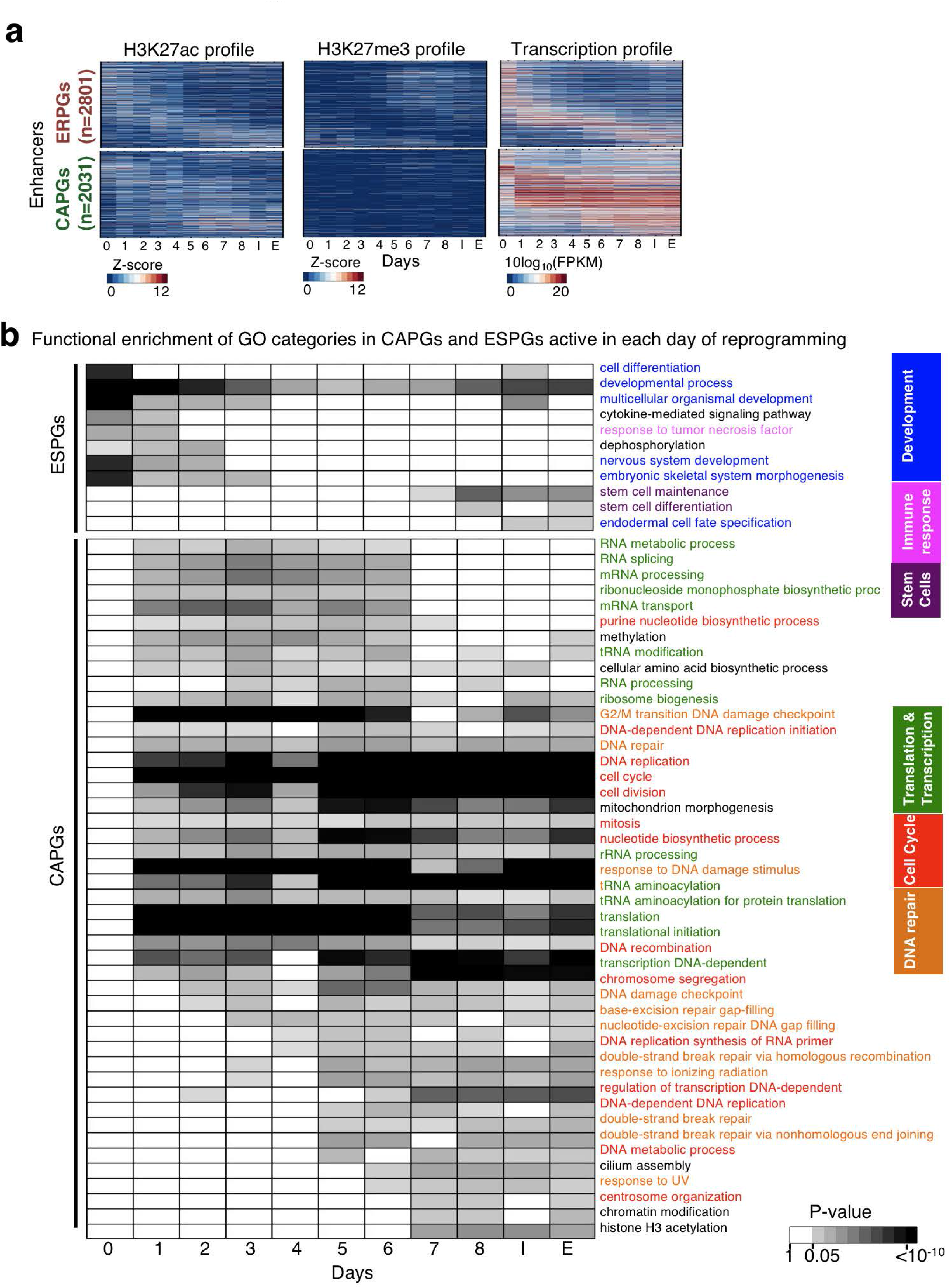
Association between epigenetic regulation (ESPGs) and cell fate genes, and between epigenetically active promoters (CAPG) and biosynthetic processes. a. Temporal profile of enhancer chromatin marks H3K27ac, H3K27me3, and gene expression of ESPGs and CAPGs that have an associated enhancer, along reprogramming. Each row corresponds to a single gene, genes are sorted according to their expression pattern and the same sorting was applied to the epigenetic marks in their corresponding enhancers. b. GO categories enriched for ESPGs and CAPGs that are active in each day of reprogramming. Gene is defined to be active in samples where RPKM is above 0.5 of the gene max value. Shades represent FDR corrected p-values (Fisher exact test). GO annotations with FDR corrected p-value < 0.01 in at least two samples are presented.

**Extended Data Fig. 12:**
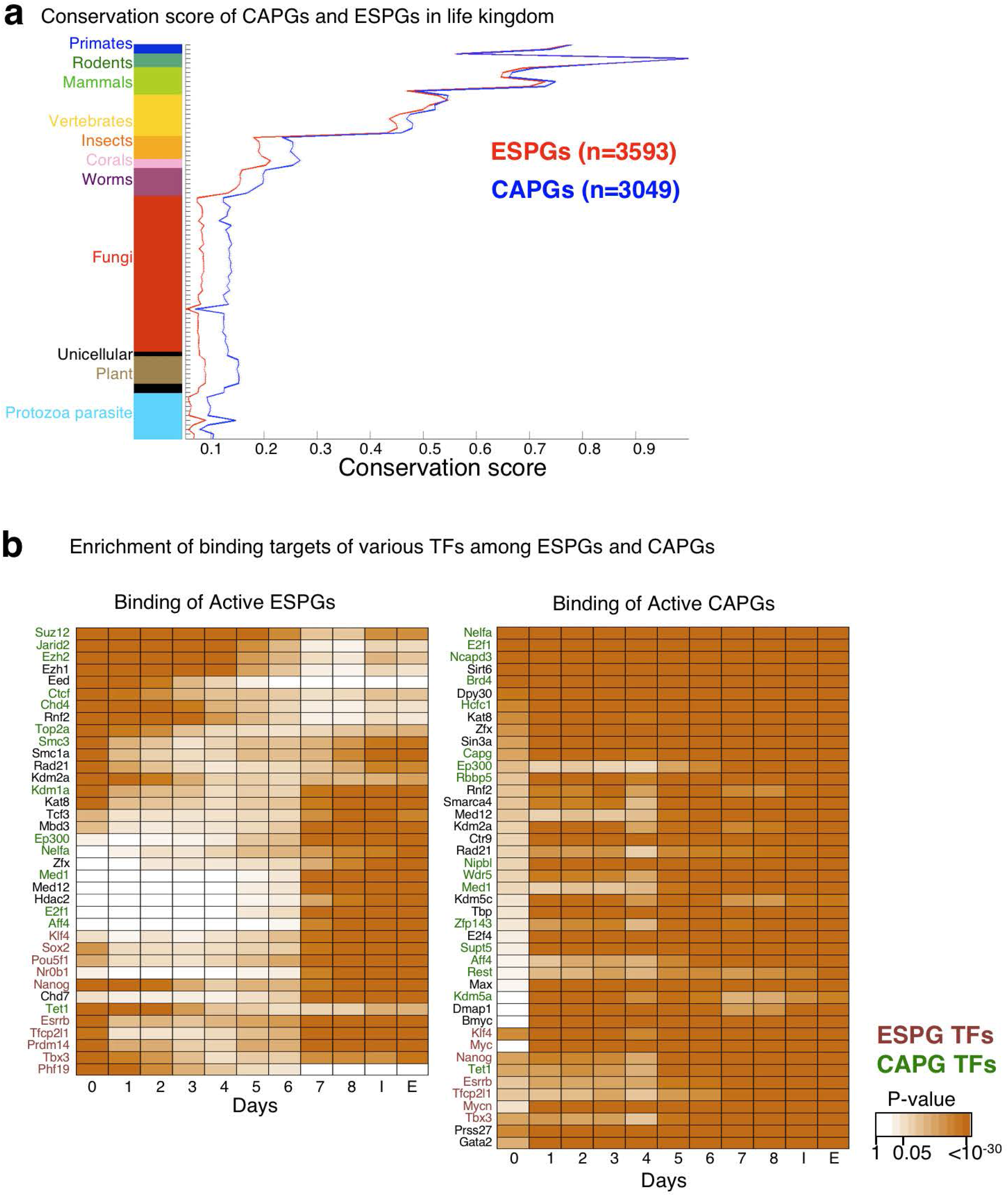
Conservation and TF binding of CAPG and ESPG groups. a. Conservation score of ESPGs (red) and CAPGs (blue), calculated with Phylogene software^74^, graph include the mean and SEM values for each gene set. Showing that CAPGs are significantly more conserved in non-vertebrate organisms than ESPGs (one tailed Wilcoxon test, p-value of nonvertebrate organisms < 10^-30^). b. Scheme showing transcription factors that significantly bind ESPGs or CAPGs that are active in each day of reprogramming, based on ChIP-seq databases^45-47^. Red– ESPGs transcription factors, Green– CAPGs factors. Orange shades– FDR corrected p-values (Fisher exact test).

**Extended Data Fig. 13:**
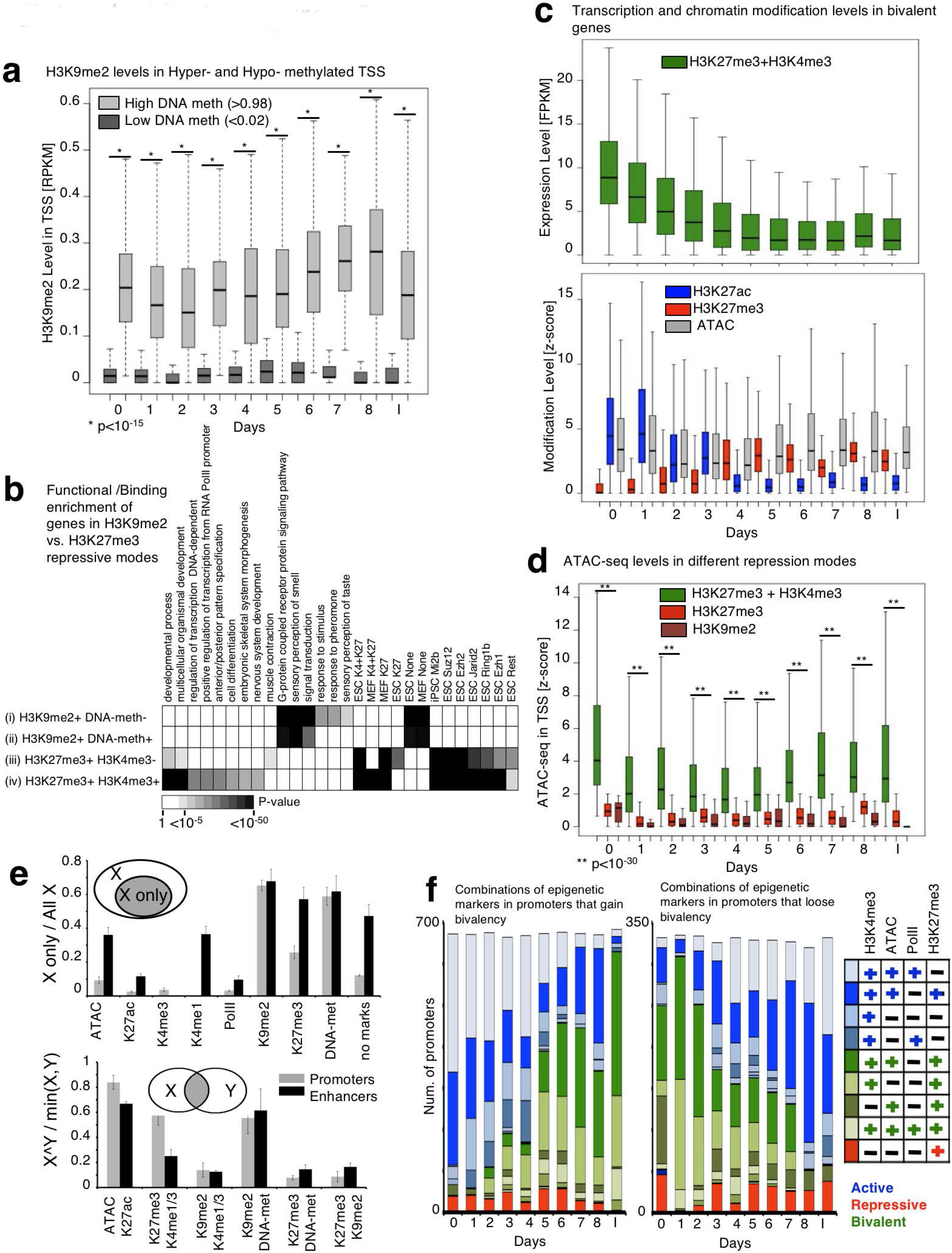
H3K9me2 and H3K27me3 show functionally distinct modes of epigenetic repression during conducive reprogramming. a. Distribution of H3K9me2 signal in highly methylated (>0.98) and lowly methylated (<0.02) gene promoters, showing association of H3K9me2 with highly methylated promoters. *p<10^-15^ (Wilcoxon test) b. Functional enrichment of genes marked by the indicated epigenetic marks in their promoters. Four gene groups were analyzed: (i) Genes with H3K9me2 without DNA-methylation (n=4159), (ii) Genes with both H3K9me2 and DNA methylation in at least two-time points (n=248), (ii) Genes with H3K27me3 without H3K4me3 (n=2220) and (iv) Genes with both H3K27me3 and H3K4me3 in at least two-time points (n=2663). Annotations include GO categories, targets of TF binding^46^, or chromatin marks measured in ESC^75^. Gray shades indicate FDR corrected Fisher exact test p-values. Annotations with corrected p-value <10^-5^ are presented. c. Top: Expression pattern of genes that gain bivalency during reprogramming (n=572). Bottom: Epigenetic modification level distribution of the same set of genes, showing a switch between H3K27ac and H3K27me3, and a constant state of accessibility. d. Chromatin accessibility of promoters marked by either repressive marks (H3K27me3, H3K9me2) or bivalent (H3K27me3+H3K4me3) modifications. Bivalent promoters are significantly more accessible than repressed promoters. **p<10^-30^ (Wilcoxon test). e. Top: Probability to observe the indicated epigenetic modification by its own in promoters (gray) and enhancers (black). Bottom: Probability to observe co-localized pairs of modifications in promoters and enhancers. f. Stacked bar chart of all combinations of the indicated chromatin modifications, as measured in promoters that gain bivalency (left) and in promoters that loose bivalency (right). Right– color code of frequent combinations (>3% of each sample).

**Extended Data Fig. 14:**
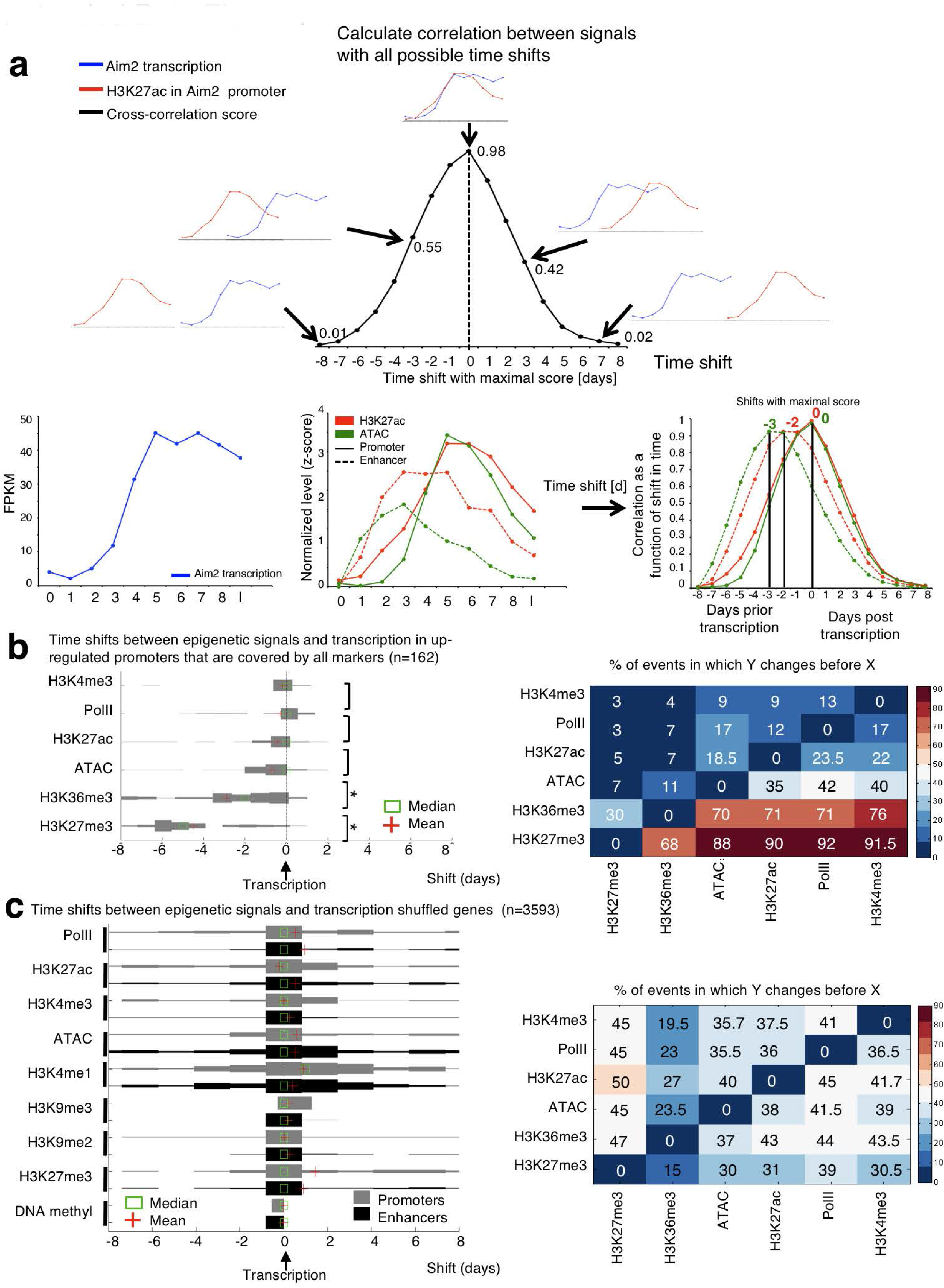
Utilizing cross-correlation method for inferring time shifts between chromatin epigenetic changes and mRNA transcription. a. Top: Cross correlation method^50,51^ measures the overlap between two signals, while shifting the signals in their x-axis (convolution). In this case, the x-axis is time. The shift that gives the highest score is defined as the temporal offset between the signals. Bottom: example for cross-correlation method calculated in the promoter and enhancer of the gene Aim2. DNA accessibility and H3K27ac signal in promoter and enhancer are shown, alongside the shift of all signals from the transcription. b. Left: Distribution of crosscorrelation temporal offset, measured between each of the indicated epigenetic modification and gene expression profile. The distribution was measured over 162 upregulated promoters that are covered by all indicated epigenetic modifications, thus comparing temporal offsets between epigenetic modifications on the same gene. Plus indicates mean, and square indicates median. *p<10^-5^ (paired-sample ttest). Right: Matrix showing the percentage of events in which the modification on the Y axis changes before the modification on the X axis. Calculated over the same set of 162 promoters as on the left panel. This matrix emphasizes that the order of events in the promoters is removal of H3K27me3, followed by deposition of H3K36me3, opening of chromatin and then deposition of other active marks (H3K27ac, PolII, H3K4me3). c. Left: Shuffling analysis was done as a negative control for cross-correlation analysis: cross correlation temporal offset was calculated between each ESPGs gene expression (n=3593) and the indicated epigenetic modification taken randomly from another gene. Showing loss of temporal order with random promoter shuffling. Right: Matrix as in (b), calculated over the shuffled promoters (n=3593), emphasizing the loss of order between the signals.

**Extended Data Fig. 15:**
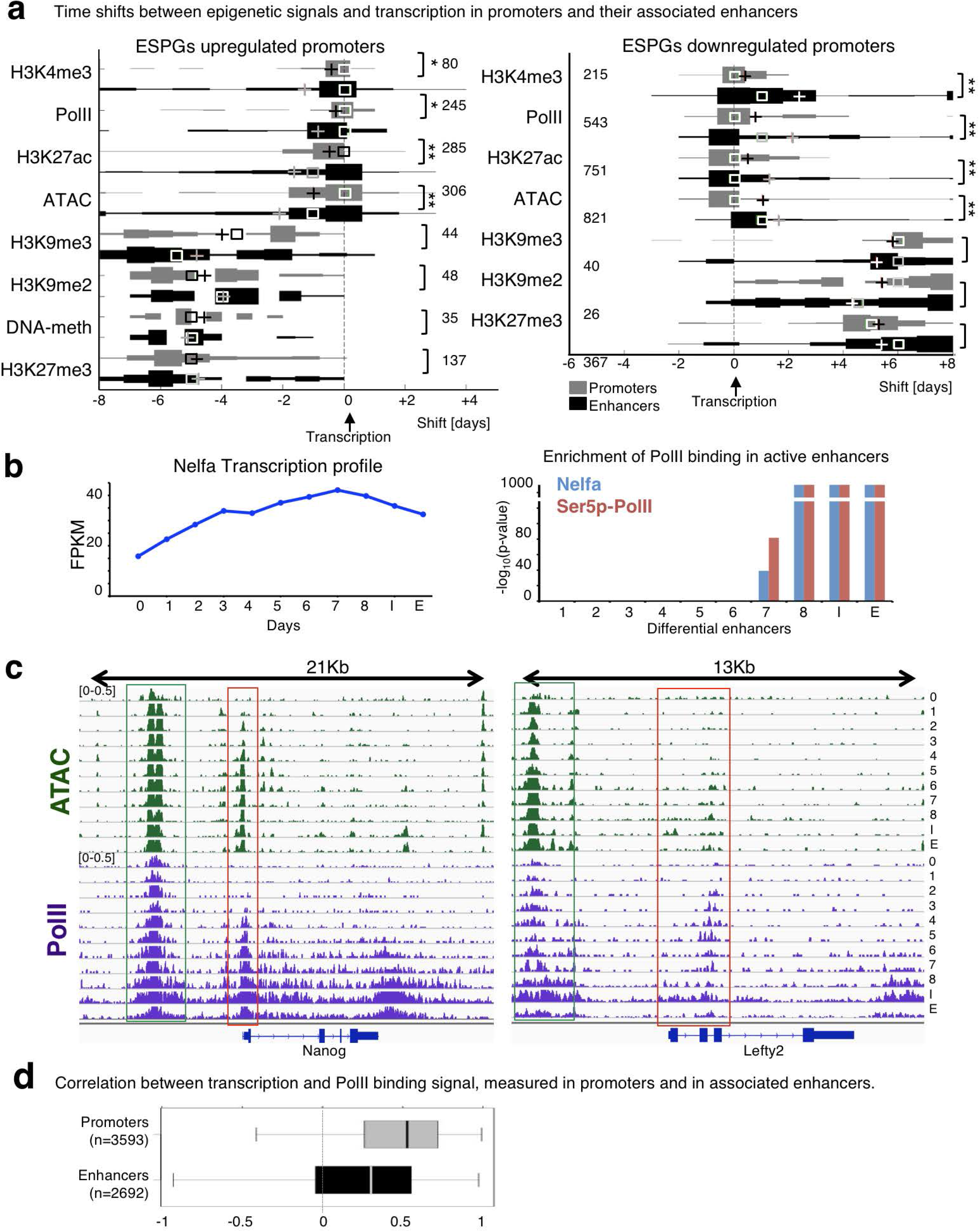
Enhancer activation and PolII enhancer binding precede activation and binding of associated promoter. a. Time shifts between the indicated chromatin modifications on promoter (gray) or associated enhancer (black) and gene expression, estimated with cross correlation method. The distribution was measured over all upregulated ESPGs (left) and downregulated ESPGs (right) which have a changing epigenetic modification (max-min z-score > 0.5) on both promoter and enhancer sites (numbers are indicated). Plus indicates mean, and square indicates median. *p<10^-2^, **p<10^-7^ (Paired-sample t-test). b. Left: Expression pattern of PolII components Nelfa during conducive iPSC reprogramming. Right: enrichment of targets bound by Nelfa and Ser5p-PolII^27^ among differential enhancers that are active in each day of reprogramming.–log_10_ of Fisher exact test p-values is indicated. c. Two cases in which PolII binds the enhancers before it binds the associated promoter. The enhancers are marked by a green box, promoters are marked by a red box. d. Correlation distribution between PolII binding signal and gene expression, as measured in promoters and in enhancers of ESPGs that are bound by PolII (numbers are indicated). Correlations were calculated for each gene, over 11 time points.

**Extended Data Fig. 16:**
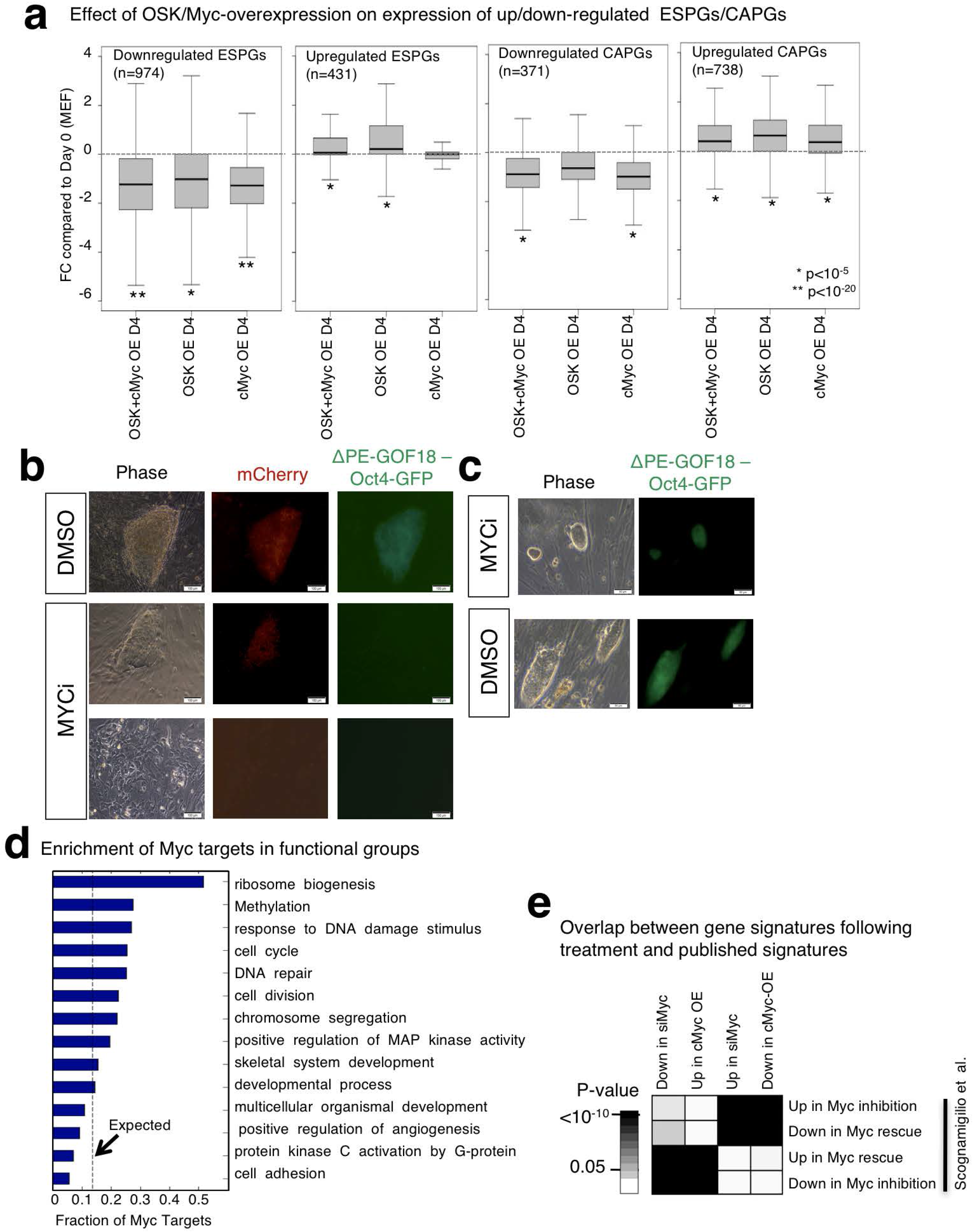
Over expression of c-Myc is sufficient for indirectly repressing somatic genes from the ESPG group. a. Distribution of Expression fold change (FC) compared to WT MEF of up/down regulated ESPGs (down regulated ESPGs are enriched for somatic genes), and CAPGs. Presented perturbations are over-expression of OSK cassette, over-expression of c-Myc cassette, or over-expression of the two cassettes together. (*p<10^-5^, **p<10^-20^, Wilcoxon test) b. Representative pictures of Mbd3^fl/-^ cells harboring mCherry-NLS and ΔPE-GOF18 Oct4-GFP cassettes after 13 days of reprogramming in the presence of MYCi. While constitutive mCherry signal was detected among growing fibroblasts or emerging colonies in MYCi conditions, we could not detect any ΔPE-GOF18-Oct4-GFP+ colonies. This indicates that Myc activity is indispensable for inducing pluripotency. Scale = 100 *μ*M. c. Representative pictures of ES cells harboring ΔPE-GOF18-Oct4-GFP reporter expanded in the presence of Myc-inhibitor. The unperturbed ΔPE-GOF18-Oct4-GFP signal indicates that MYCi is dispensable for maintaining pluripotency as recently shown^55^. Scale = 100 *μ*M. d. Fraction of Myc targets in significantly induced and repressed GO categories, compared to what is expected by random (dashed line). e. Overlap between differential genes detected in Myc perturbation experiments, and differential genes detected in previous published perturbations^55^, showing consistency between the two experimental settings in two different labs. Fisher exact test p-values are presented.

**Extended Data Fig. 17:**
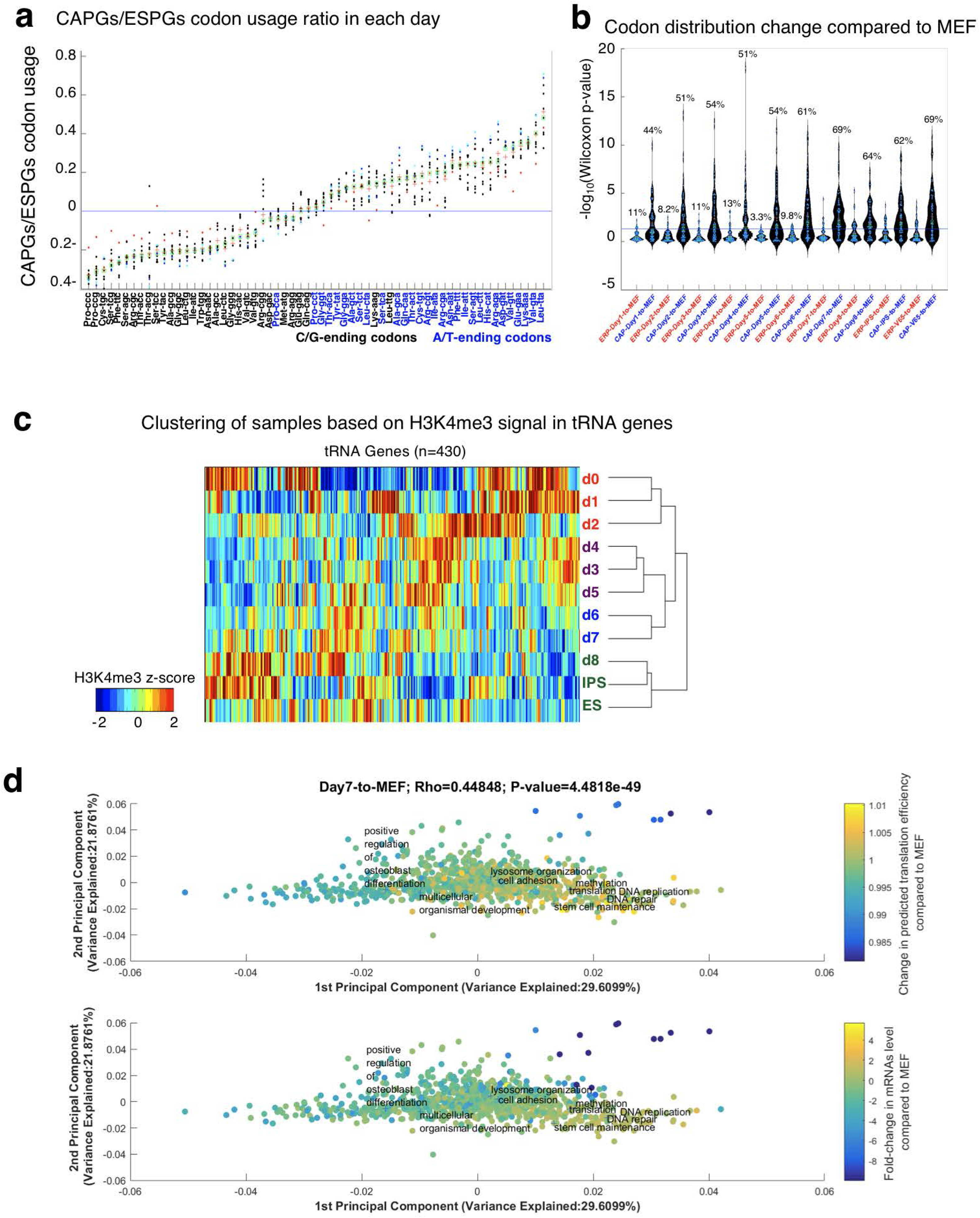
Distinct and rapid change in codon usage is specific to CAPGs. a. ESPGs prefer G/C-ending codons, while CAPGs use A/T-ending ones. Scatter plot analysis of the CAPGs-to-ESPGs ratio of codon usage. For each codon, shown are 11 calculated CAPGs-to-ESPGs ratios of codon usage along reprogramming and in ESCs. The mean and median values of each codon are shown as red crosses and green squares, respectively. Three time points are color-coded: red = CAPGs-to-ESPGs ratio in MEF; blue = CAPGs-to-ESPGs ratio in iPS; light blue = CAPGs-to-ESPGs ratio in ES. b. CAPGs change significantly their codon usage compared to MEF already in day 1 of reprogramming, where ESPGs are much less variable. Violin plot of the significance (-log_10_(Wilcoxon p-value)) of the differences between the usage of each of the 61 sense codons in each day compared to MEF, calculated for ESPGs (red) and CAPGs (blue) sets (for each day separately). c. Hierarchical Clustering of various time points/cell types based on the H3K4me3 signature in the vicinity of individual tRNA genes. (n=430, Spearman correlation metric and average linkage were used). d. Projection of the tRNA translation efficiency and mRNA expression changes on the Codon-Usage Map. PCA projection of GO category gene-sets. The location of each gene set in this space is determined by the average codon usage of all the genes that belong to it. The % variance spanned by the first and second PCs is indicated on the x and y axis, respectively. The color code represents the predicted translation efficiency of tRNAs and expression change of mRNAs (upper and lower panels, respectively) in day1 compared to MEF. Top: each gene category is color coded according to the relative change in the availability of the tRNAs that correspond to the codon usage of its constituent genes, averaged over all genes in the category; the tRNA availability of each individual gene was calculated similarly to the tAI measure of translation efficiency, where the expression of individual tRNA genes was evaluated by the H3K4me3 reads in its vicinity. A red color for a given gene category indicates that on average the genes in that category have codons that mainly correspond to the tRNAs that are induced in day1, whereas a blue color indicates that the codon usage in the categories is biased toward the tRNAs that were repressed in day1. Bottom: Changes at the mRNA level, averaged over all the genes in each gene category as in the upper panel, where here too red means that the genes were induced in day1.

**Extended Data Fig. 18:**
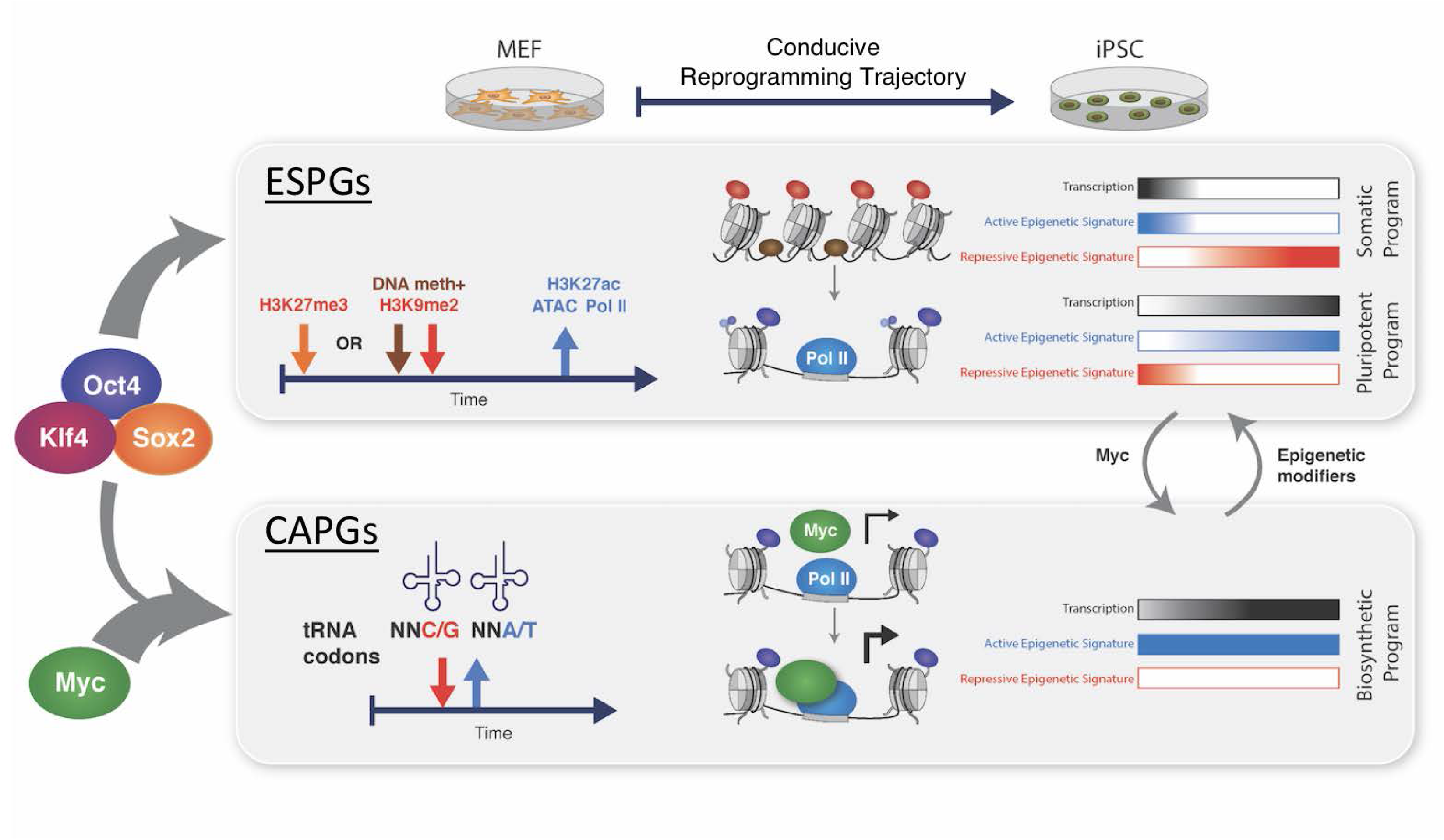
Schematic model for molecular drivers and changes underlying conducive reprogramming to murine ground state pluripotency. Interleaved epigenetic and biosynthetic reconfigurations rapidly commission and propel conducive reprogramming toward naïve pluripotency. The latter requires the coordinated activity of two gene groups, CAPGs and ESPGs, each with distinct modes and kinetics of regulation.

## Supplementary Tables Figure Legends

**Supplementary Table 1: Summary of sequencing Metadata.** Number of aligned reads, number of peaks, and sequencing protocol of each library, for Mbd3^f/-^, Gatad2a^-/-^, WT-1 and WT-2 engineered mouse reprogrammable systems.

**Supplementary Table 2: Gene expression values of 8042 differential genes in four different drug inducible reprogramming systems.** Calculated from PolyA+ RNA-seq, including protein-coding genes, pseudogenes, and lncRNAs, measured in the four different secondary reprogramming systems: Mbd3^f/-^, Gatad2a^-/-^, WT-1 and WT-2. Values are unit normalized FPKM (see Methods).

**Supplementary Table 3: Gene expression values of 8705 differential genes in Mbd3^f/-^ system.** Gene expression values of 8705 differential genes in Mbd3^f/-^ system, including polyA+ genes (protein-coding, pseudogenes, lncRNAs) and small RNA-seq (rRNA, miRNA, snoRNA). Values reported are FPKM.

**Supplementary Table 4: Differential lncRNAs during conducive naïve iPSC reprogramming trajectory.** Expanded lncRNA analysis beyond Ensemble annotation utilizing a PLAR-generated lncRNA dataset (see Methods). 560 differential lncRNA were identified during reprogramming, out of them 221 differential lncRNA not annotated in Ensembl.

**Supplementary Table 5: Clusters of enhancers with different demethylation dynamics.** Datasheet annotates clusters of enhancers with different demethylation dynamics as presented in Fig. 3b. Cluster 8 includes enhancers and super-enhancers with fast demethylation.

**Supplementary Table 6: ESPG and CAPG differential groups.** Detected differential gene groups during conducive iPSC reprogramming, including all differential genes and the differential genes subsets annotated as genes with epigenetically switched promoters (ESPGs) and genes with constitutively active promoters (CAPGs). Gene list is further divided to up-regulated and down-regulated list of each gene group.

